# Repair of DNA double-strand breaks leaves heritable impairment to genome function

**DOI:** 10.1101/2023.08.29.555258

**Authors:** Susanne Bantele, Irene Mordini, Alva Biran, Nicolas Alcaraz, Alice Wenger, Nils Krietenstein, Anja Groth, Jiri Lukas

## Abstract

Upon DNA breakage, a genomic locus undergoes alterations in 3-D chromatin architecture to facilitate signaling and repair. While cells possess mechanisms to repair damaged DNA, it is unknown whether the surrounding chromatin is restored to its naïve state. We show that a single DNA double-strand break (DSB) within a topologically-associated domain (TAD) harboring conformation-sensitive genes causes lasting chromatin alterations, which persist after completion of DNA repair and feature structural changes, chromatin compaction and loss of local RNA species. Unexpectedly, these newly-acquired features of post-repair chromatin are transmitted to daughter cells and manifest as heritable impairments of gene expression. These findings uncover a hitherto concealed dimension of DNA breakage, which we term post-repair chromatin fatigue, and which confers heritable impairment of gene function beyond DNA repair.

## Introduction

The central function of a cell’s genome is not only to hold the genetic information that is read by the transcription machinery, but also to sustain heritable patterns of its activity and ability to respond to endogenous and exogenous cues that direct cell-fate decisions. Four key factors cooperate to preserve memory of gene activity: DNA sequence, DNA and histone modifications, associated non-coding RNAs, and the overarching three-dimensional (3-D) chromatin structure. The latter is established by hierarchical folding at multiple scales starting from small loops, followed by larger topologically associated domains (TADs) and higher-order chromatin compartments, which ultimately make up structural chromosome territories in cis and functional territories (hetero-and euchromatin or B-and A-compartments) in trans (*1–5*).

Due to its ability to affect the proximity of linear genomic segments, chromatin folding has been widely assumed to impact transcription efficiency and dynamics. While this is clearly the case in some specific tissues or developmental contexts, the causative role of 3D genome organization (and its heritability across successive cell generations) for gene activity is still being discussed. On one hand, depletion of topological insulators only causes mild gene deregulation (*6*, *7*). On the other hand, mounting evidence indicates that specific gene regulation upon cellular stimulation (*8–11*), differentiation (*12–19*), oncogenic transformation (*12*, *20–22*) or during evolution can be modulated by local topology modifications capable of aligning, disrupting or even re-programming promoter-enhancer interactions and accommodating regulatory factors. Thus, rather than a static picture of ‘pre-determined’ 3-D chromatin context for gene expression, a much more dynamic model is emerging where the topological plasticity of the genome is strategically leveraged when cells need to quickly adapt to a new environment or respond to stimuli including various stress assaults.

An outstanding form of stress that can impair 3-D genome organization and derail cell identity include DNA double-strand breaks (DSBs). This is because interruption of DNA integrity provokes enzymatic responses within large segments of the surrounding chromatin, which can extend up to several megabases away from the primary DNA lesions. By analogy to a transcriptional start site, which must accommodate bulky protein complexes and potentially interact with enhancer regions to be transcribed, a genomic locus harboring DNA lesions must be accessed by DNA-end-processing nucleases, signaling proteins, DNA repair complexes, and in case of recombination-based repair undergo directional movement to align with a homologous repair template. It is therefore not surprising that besides acute changes in chromatin compaction and mobility, post-translational modifications of histones and the DNA repair process per se, extensive topological rearrangements including chromatin compartmentalization are also among inevitable consequences of DNA breakage (*23–29*). The pre-existing chromatin context defined by boundary factors of chromatin organization may facilitate the formation of such DSB-induced chromatin domains to increase the efficiency and foster the fidelity of DNA repair (*24*, *30*, *31*).

From an evolutionary perspective, the key task of a cell is to repair damaged DNA within the same cell cycle before the cell separates chromosomes in mitosis and transmits parental genomes to daughter cells. This is achieved through DNA damage checkpoints, which transiently halt cell cycle progression to generate time for DNA repair per se but also for coordinating local repair reactions with gene expression activity in the affected genomic loci. Indeed, DSBs in active chromatin induce transient suppression of local transcription to prevent production of faulty transcripts, which could otherwise cause potentially harmful collisions between DNA and RNA transactions (*32–34*). The aim of such coordinated regulation is to restore integrity of DNA sequence and thus allow recovery of a supposedly functional genomic locus to be passed on to the daughter cells. It has been well established that compromised DNA repair fidelity that can lead to mutations in gene-coding sequences or their regulatory elements undermines this process and can lead to devastating cell-fate consequences including developmental defects and oncogenic transformation. However, it remains unknown whether the extensive changes in three-dimensional topological re-arrangements that are inevitably coupled to DSB signaling and repair are also fully restored back to the naïve state. This is an important gap in our knowledge of physiological consequences of DNA breakage because heritable changes in higher-order chromatin arrangement could, in principle, undermine gene activities and thus derail functional robustness of progenies of cells despite the integrity of primary DNA sequence is restored after a DSB insult.

Here, we address this issue by systematically dissecting the lasting consequences of DSB repair on 3-D chromatin arrangement and the associated gene expression. Using single, Cas9-induced DNA double-strand breaks directed to specific locations withing an entire TAD harbouring both protein-coding genes and regulatory RNA species, we demonstrate that a repaired genomic locus is not able to accurately resume regular transcription, even when the lesion has been generated (and subsequently repaired) in megabase distances from the gene itself. These defects coincide with an altered topological makeup of the affected locus and are inherited to the next generations of daughter cells. Hence, we propose that a DSB insult to genome integrity has a lasting impact on cellular physiology that reaches beyond the mutagenesis of the primary DNA sequence.

## Results

### A single Cas9 DSB anywhere in the c-MYC TAD has a protracted effect on c-MYC protein expression

To start dissecting coordination between the repair of damaged DNA and topological configuration of the neighboring genomic locus, we asked whether a single Cas9 DNA double-strand break affects transcriptional activity of genes located outside of the actual breaksite. As an experimental system, we chose the c-MYC locus, which is embedded in the more than 3 megabases large TAD on chromosome 8 (hereafter called the c-MYC TAD, Fig. 1A and Fig. S1A). The c-MYC gene itself was previously shown to be transcribed in a topology-dependent manner (*35*, *36*), making the c-MYC TAD an ideal environment to quantitatively interrogate the dynamics, magnitude, and temporal fluctuation of gene expression after disrupting 3-D chromatin arrangement by DNA breakage. We established twelve Cas9 cutsites spanning the entire c-MYC TAD (Fig. 1A, scissor symbols), including two cutsites within the c-MYC open reading frame (ORF) as a positive control and confirmed specific cutting using the T7E1 surveyor assay (Fig. S1B). Cutting efficiency was inferred on the single cell level measuring 53BP1 accumulation at DNA FISH-labeled c-MYC loci, where the represented T7E1 cleavage (Fig. S1B; MYC-ORF2) correlated with up to 70% of 53BP1-positive Cas9 MYC-ORF2 loci (Fig. 1C top, 6h timepoint). Since Cas9 cutting and subsequent DNA repair are dynamic processes, such steady-state levels of positive cutting indicate efficient cleavage. Cas9 was delivered to cells as in vitro assembled ribonucleoprotein complex (RNP) with the respective guide RNA to ensure fast cutting without extended expression of Cas9 in the cells. Using this system, we first determined the acute transcriptional response of the c-MYC gene upon a single Cas9 cut in the c-MYC TAD using single cell quantitative image-based cytometry (QIBC; Fig. 1B). 53BP1 accumulation at Cas9 cutsites was used as a measure of DNA repair kinetics, which peaked around six hours after Cas9 RNP transfection with up to 70% of cutsites bound by 53BP1. After 24h, the repair response was largely completed with roughly 80% of cutsites free of 53BP1 association (Fig. 1C, top).

**Fig. 1.**
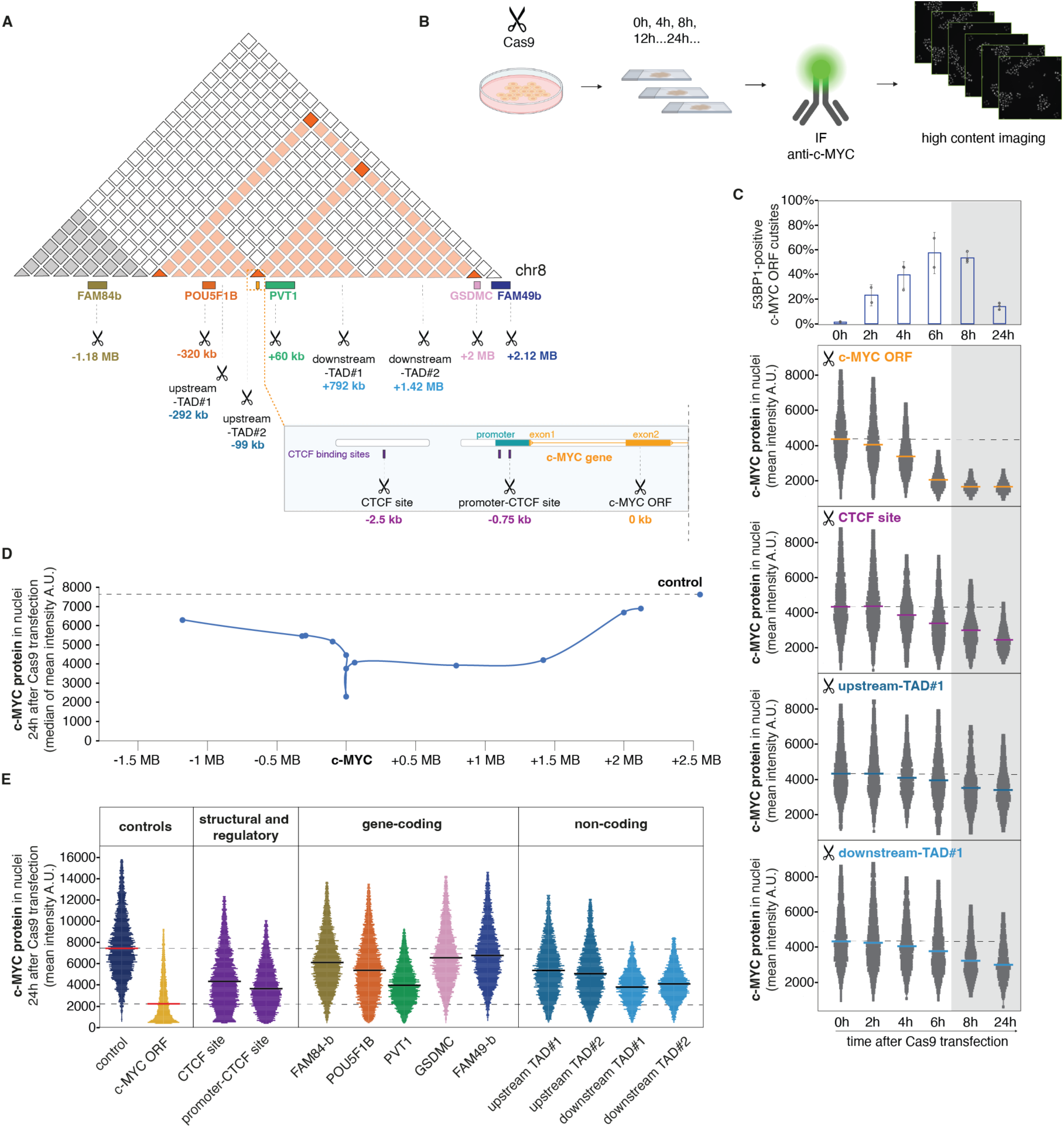
A Single Cas9 DNA DSB within the c-MYC topological domain has a protracted effect on c-MYC protein expression. (A) Schematic depiction of the c-MYC TAD on chromosome 8 (highlighted in orange, see also figure S1A) with indicated Cas9 cutsites (scissor symbols). The inset magnifies the c-MYC coding region including upstream elements with CTCF binding sites. Numerical values indicate the genomic distance of a given Cas9 cutsite from the c-MYC coding region. Colored boxes indicate locations of transcripts encoded in the c-MYC TAD. (B) Schematic representation of a quantitative image-based cytometry (QIBC) experiment to quantify protein expression of c-MYC in single cells by high content imaging. In a typical experiment, HeLa cells (unless indicated otherwise) were subjected to a single Cas9 cut by transfection of in vitro assembled ribonucleo-protein complexes (Cas9 RNPs), fixed at indicated timepoints, stained for the c-MYC protein by immunofluorescence, and analyzed by QIBC. (C) Temporal alignment of DNA repair kinetics as measured by immunostaining of 53BP1 at c-MYC ORF Cas9 cutsite marked by DNA FISH (average of 2-3 biological replicates, n=120-200 c-MYC FISH foci per timepoint per replicate, error bars show the standard deviation) with mean c-MYC protein levels per cell measured by QIBC after Cas9 cutting at indicated sites and timepoints (n=1500 cells per timepoint, horizontal bars indicate the median of the mean c-MYC protein level per cell, dotted lines represent the mean value of c-MYC protein expression before Cas9 RNP transfection). (D) Mean c-MYC protein expression at indicated genomic distances from the c-MYC ORF 24h after Cas9 RNP transfection; n=7000 cells per RNP, plot shows values from one representative of at least three or more biological replicates (see also figure S1B-G). (E) One representative biological replicate from the same experiment as in (D). Horizontal bars indicate the median of the mean values; (n=7000 cells per each depicted condition). Colors throughout the figure match the schematic depiction in (A).

As expected, the c-MYC protein expression gradually decreased within the first six hours after the Cas9 transfection in the c-MYC ORF (Fig. 1C). Unexpectedly, Cas9 cuts outside of the c-MYC ORF also decreased c-MYC protein expression during this acute repair phase, even when located as far as 300 kb upstream (upstream-TAD#1) or 800 kb downstream (downstream-TAD#1) of the c-MYC ORF (Fig. 1C). While the decrease of c-MYC protein after cutting at remote locations was initially weaker compared to cutting directly in the c-MYC ORF, it persisted and became gradually even more pronounced at timepoints when 53BP1 largely dissociated from the chromatin near the cutsite (Fig. 1C). Thus, the reduced c-MYC protein expression persists beyond the repair of the actual DSB lesions, even if these are located outside of the coding region within the same TAD. This effect was neither due to the R-loop formed at the Cas9 cleavage site, nor the DNA-binding of Cas9 per se, as catalytically dead Cas9 did not affect c-MYC expression even after 24h (Fig. S1D). Similarly, a single nick by mutated Cas9 was not sufficient to repress c-MYC protein expression independent of which strand was cleaved, even when it was in the c-MYC ORF itself (Fig. S1E). Consistent with the single-time-point analysis, live-imaging of endogenously tagged c-MYC-AID-mEGFP HeLa cells (Fig. S1Fi-iii) confirmed protracted c-MYC repression in the entire population of hundreds of cells after Cas9-generated DSBs either within or outside the c-MYC ORF (Fig. S1G, bottom). Importantly, this effect was not due to DSB-associated effects on cell proliferation, which was unhampered in cells with only a single Cas9 cut (Fig. S1G, top).

We therefore wondered whether the extent of c-MYC repression would correlate with the distance between the breaksite and the c-MYC coding sequence. To this end, we quantified c-MYC protein levels 24h after a single Cas9 cut at each of the twelve sites within the c-MYC TAD, spanning between 1.5 MB upstream and 2.5 MB downstream of the c-MYC ORF (Fig. 1A). All cells affected by a Cas9 cut showed reduced c-MYC protein levels compared to control cells transfected with non-targeting Cas9 (termed “control” hereafter, marked by dotted line), with the strongest repression within or adjacent to the c-MYC ORF. Importantly, while the extent of c-MYC repression gradually decreased with the distance from the c-MYC gene, it remained significantly below the control levels even after cuts at the extreme c-MYC TAD boundaries (Fig. 1D, S1C). Interestingly, it did not matter whether the cutsite was placed within a transcribed region of coding or non-coding RNAs across the c-MYC locus, structural elements like CTCF sites or the promoter of the c-MYC gene, since in all cases a partial c-MYC gene repression was observed (Fig. 1E). Taken together, these results suggest that a DSB at any targeted position within the topologically defined environment of the c-MYC gene partially dampens c-MYC protein production even after DNA repair is finished.

### A single Cas9 DNA DSB affects the transcriptional output across the entire c-MYC TAD

Given the fact that c-MYC itself has been described as a topology-sensitive gene (*35*, *36*), we next asked whether the observed expression dampening by remote DSBs is confined to the c-MYC TAD and whether it is exclusive to the c-MYC gene. Furthermore, we wanted to know whether the effects observed by quantifying protein levels originate from the impairment of gene expression. We chose a subset of Cas9 cutsites (Fig. 2A) and assayed by RT-qPCR acute (6h) and lasting (24h) RNA production of c-MYC, POU5F1B and FAM84B genes located within or in immediate neighborhood to the c-MYC TAD (Fig. 2A). Expression of all three mRNAs (Fig. 2B i-iii) was partially decreased after 6h of any of the Cas9 cuts, confirming a wide reach of acute transcriptional repression during ongoing DSB repair. Analogous to the protein levels (Fig. 1C), c-MYC mRNA remained repressed also after 24h (i.e., after completion of the bulk of DNA repair) at all cutsites outside the c-MYC ORF (Fig. 2B i). It is noteworthy that the c-MYC mRNA was immediately repressed to full extent after 6h when the cut was placed in the c-MYC ORF, while all other cuts caused a gradual reduction of c-MYC transcripts. The POU5F1B mRNA (Fig. 2B ii) transcribed from a locus upstream of the c-MYC ORF also underwent gradual and lasting repression by cutting at all sites upstream of the c-MYC ORF, which is located adjacent to several binding sites of a regulator of chromatin organization, CTCF. A cut downstream of these CTCF sites (downstream-TAD#1) did not affect POU5F1B expression, hinting at a potential influence of topological boundary elements, at least for the expression of some genes.

**Fig. 2.**
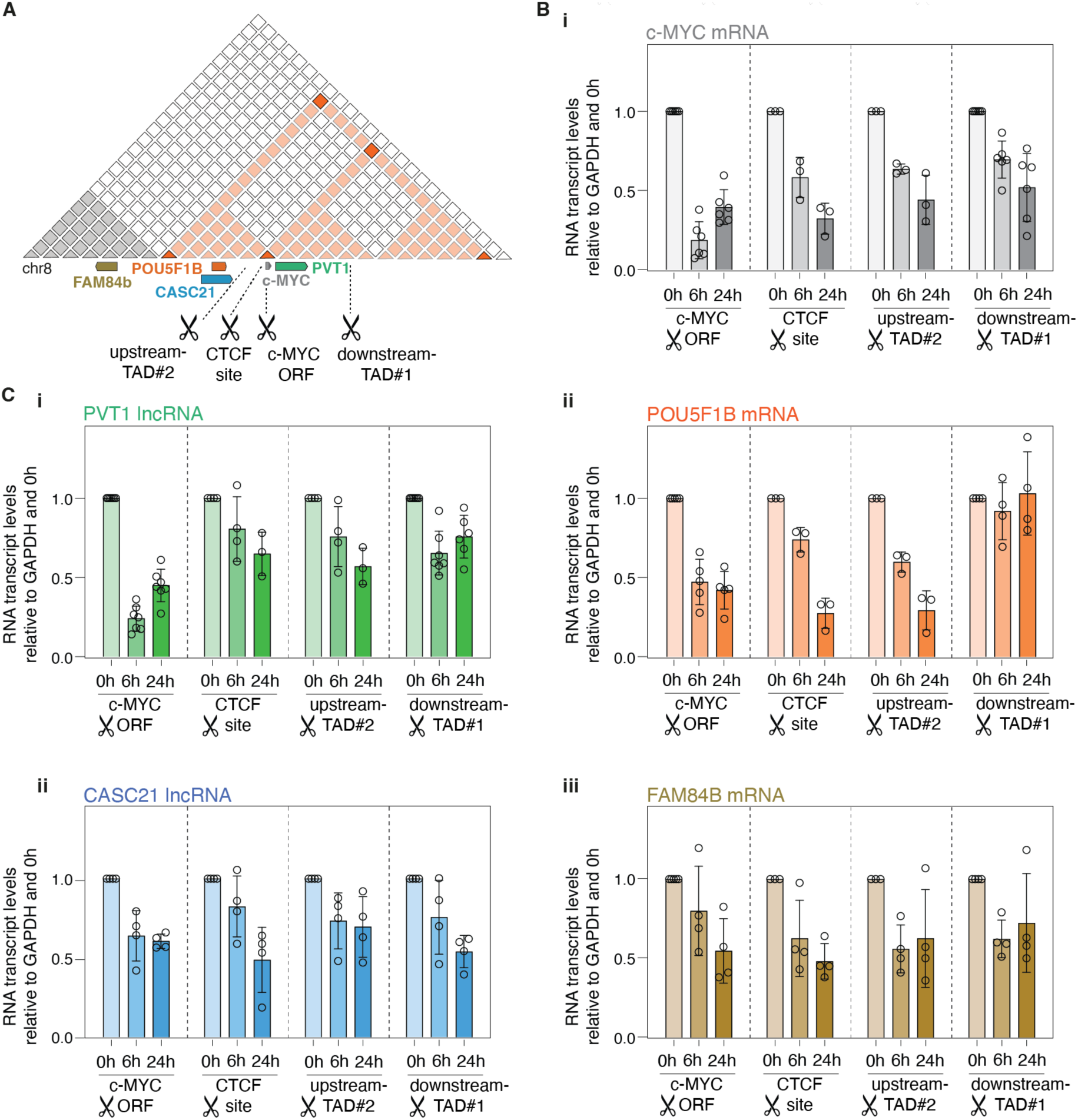
A Single Cas9 DNA DSB affects the transcriptional output across the entire c-MYC TAD. (A) Schematic depiction of the c-MYC TAD (highlighted in orange) with indicated subset of Cas9 cutsites (scissor symbols). Colored boxes mark genomic locations of mRNA and lncRNA transcripts quantified in this figure. (B)(i-iii) mRNA transcript levels of c-MYC (i), POU5F1B (ii) and FAM84B (iii) at indicated timepoints after Cas9 RNP transfection measured by RT-qPCR; Cas9 cutsites are indicated below each panel. All values are plotted relative to GAPDH and normalized to the 0h timepoint using the ΔΔCT method; bars show the average of n=3-6 replicates per condition with 3 technical replicates each, circles show biological replicates. (C)(i-ii) Same analysis as in (B) for PVT1 (i) and CASC21 (ii) lncRNA transcript levels as indicated.

Interestingly, expression of PVT1 and CASC21 long non-coding RNAs derived from loci down-and up-stream of the c-MYC ORF, respectively (Fig. 2A) were also affected by Cas9 cuts outside their coding sequences with increasing repression from acute to late timepoints (Fig. 2C i-ii). The only exception from this temporal pattern was the pronounced repression of the PVT1 lncRNA already after 6h upon Cas9 cutting in the c-MYC ORF (Fig. 2C i). This might either be due to the proximity of the PVT1 and c-MYC genes, or due to interactions between the c-MYC mRNA and the PVT1 lncRNA (*37*, *38*). Importantly, transcript repression was not due to low c-MYC levels, as depletion of c-MYC by siRNA did not reduce transcription in adjacent genes (Fig. S2A, B) Collectively, these experiments support and extend the emerging picture of a long-range and lasting impairment of gene expression within a locus bearing a repaired DSB, with potential influence of topological boundaries on insulating the neighboring loci from this effect.

### Alteration of c-MYC expression after a single Cas9 DSB is heritable

Re-formation of the interphase chromatin organization upon entry in the G1 phase provides an opportunity for the cell to reset the chromatin state and reverse potential alterations in gene expression (*39*, *40*). We therefore asked whether daughter cells can resume normal transcriptional activity in the locus wherein their mother incurred, and repaired, a DSB. To this end, we chose six Cas9 cutsites across the c-MYC TAD (Fig. 3A) and assayed c-MYC protein production using QIBC after 24h, 48h and up to 96h after Cas9 RNP transfection. Strikingly, the extent of c-MYC repression after each cut measured at the 24h timepoint was maintained also in the later time points, where the bulk of cells repaired the Cas9-inflicted DSBs and continued to divide without detectable cell cycle alterations (Fig. 3B, S3A, B). As a control for cell cycle arrest, we introduced Cas9-mediated cuts at telomeric repeats of all chromosomes (Fig. S3A, B); these cells remained arrested in G2 and only resumed natural cell cycle distribution after 96h (Fig. S3A; red arrow; B; red boxes). Noteworthy, such cell cycle arrest per se did not affect c-MYC protein production in subsequent cell generations, indicating that reduction of gene expression after DSBs is locally confined to the TAD bearing the DNA lesion and not due to global stress signaling. In support of this conclusion, RT-qPCR confirmed partial suppression of other transcripts within the c-MYC TAD, which was sustained up to 72 h after Cas9 RNP transfection at all tested sites (Fig. 3C). Thus, a single DSB seems sufficient to cause heritable impairment of gene expression in a large, topology-defined region surrounding the repaired DNA lesion, providing the first indication of a memory of DNA damage that can be propagated to next generations of cells and that goes beyond the restoration of DNA sequence integrity.

**Fig. 3.**
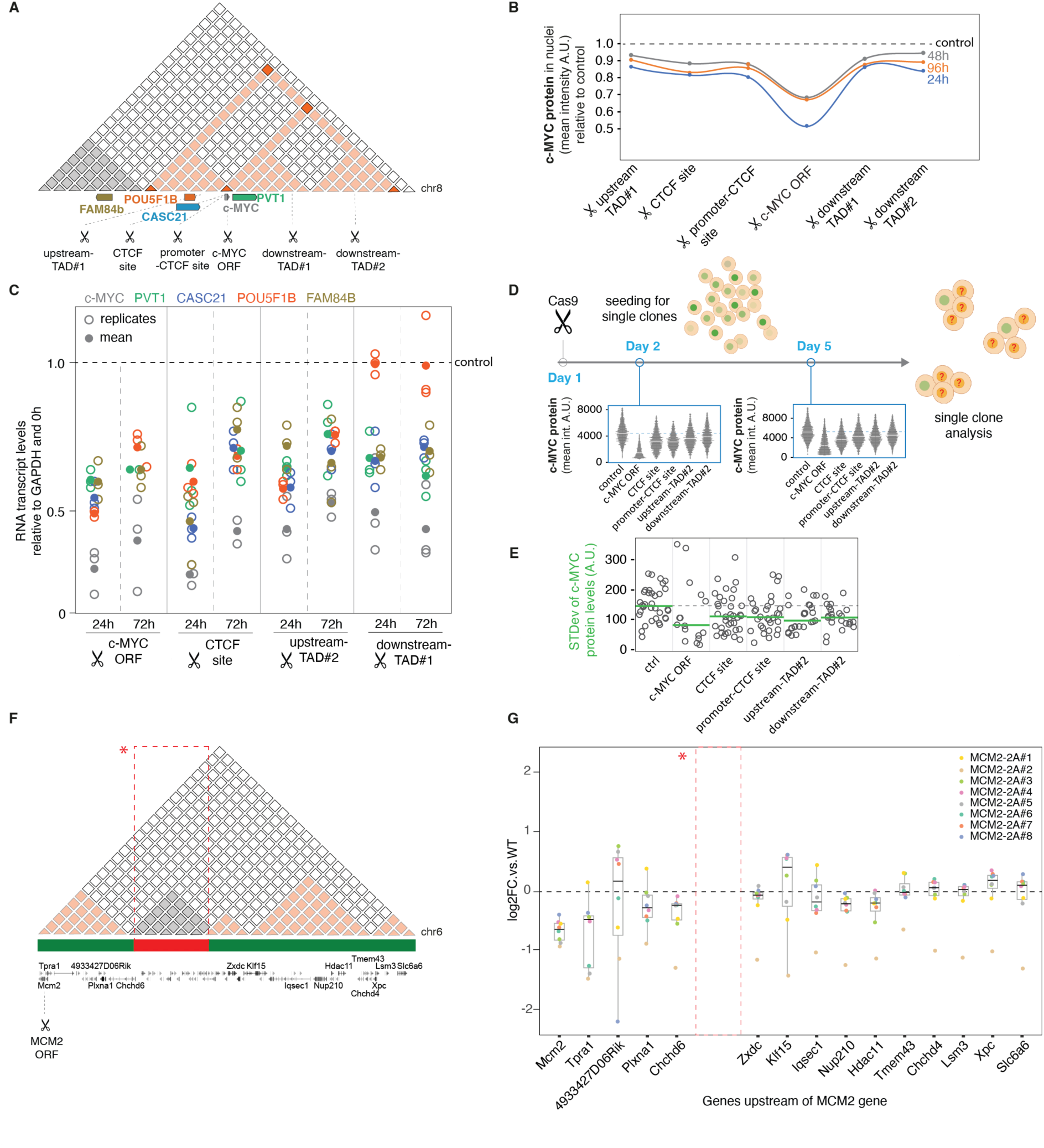
Alteration of c-MYC expression after repair of a single Cas9 DSB is heritable. (A) Schematic depiction of the c-MYC TAD on chromosome 8 (highlighted in orange) with indicated Cas9 cutsites (scissor symbols). (B) Mean c-MYC protein expression at indicated timepoints after Cas9 cutting at indicated cutsites (see also figure S3A-B). Values are plotted relative to the non-targeting Cas9 control sample at the respective timepoint and represent the median of the mean protein levels per cell; n=13000 cells per sample. (C) Relative mRNA and lncRNA transcript levels (obtained and calculated as in Fig. 2) measured by RT-qPCR at indicated timepoints and Cas9 cutsites (3 biological and 3 technical replicates). Empty dots represent individual replicates; filled dots represent the mean. (D) Schematic representation of single clone expansion experiment. Cells were subjected to a single Cas9 cut, allowed to repair for 24h, seeded for single clones and allowed to expand through at least two subsequent cell divisions (>4 cell stage). Clones were then analyzed for mean c-MYC protein expression per cell as in Fig. 1. Plots represent the mean expression of c-MYC protein per cell; n=2000 cells for each depicted condition; horizontal bars mark the median of the mean. (E) Variance of mean c-MYC protein expression within single clones (see also figure S3C); horizontal bars mark the median of the mean; dotted line shows the non-targeting Cas9 control sample; n=16-35 clones per condition. (F) Schematic depiction of the MCM2 genomic environment on mouse chromosome 6 (A compartments labeled by green bars, B compartment labeled by red bar and highlighted by red box, see also figure S3D) with indicated Cas9 cutsites (scissor symbols). (G) Expression of genes in upstream proximity to the *Mcm2* gene after genome editing in mouse embryonic stem cells (mESCs), see also figure S3E. Plot shows relative expression of indicated, differentially regulated genes (|log2 FC| > 0.58, adjusted P-value < 0.01) in 8 analyzed MCM2-2A clones and one rescue cell line (MCM2-R#2). MCM2-WT was used to normalize expression levels of the depicted genes, n = 20 (8 clones in 2-5 biological replicates), for MCM2-2A mutant, n=22 (8 clones in 2-8 biological replicates). Red box with asterisk indicates the location of the intersecting B compartment (derived from Micro-C data (4DNFILGE5LQU, 4N2 portal)(*41*)) plotted in Fig. S3D.

To directly test this conclusion, we assessed whether the heritable transcriptional attenuation remains over several successive cell generations at clonal level. We inflicted single Cas9 cuts and let cells repair the lesions for 24h. We then isolated single cells and let them clonally expand until they reached at least the four-cell stage after two rounds of mitosis (Fig. 3D). Consistent with the prediction derived from our previous data, QIBC analysis confirmed persisting c-MYC protein reduction throughout successive proliferation cycles (Fig. 3B, D). Strikingly, within each analyzed single clone, we not only recorded the same pattern of partial decrease of the mean expression level of c-MYC, but also a consistent decrease of the variance of c-MYC expression compared to control cells (Fig. 3E; Fig. S3C). Hence, the clonally expanded cells that recovered from a DSB within the c-MYC TAD displayed a reduced dynamic range of c-MYC expression along with overall lower levels of c-MYC protein expression.

Importantly, by mining the precious resource of cellular clones derived from mouse embryonic stem cells after DSB-based editing of the *Mcm2* gene (*42*), we found that heritable changes in transcription post DNA repair were not restricted to the c-MYC TAD or the HeLa cell line. Specifically, RNA-seq analysis revealed that the expression of genes in upstream proximity to the edited MCM2 sequence (up to 700-800 kb, Fig. 3F-G, S3D-E) was deregulated in several independent clones. This deregulation could not be rescued by reversion of the MCM2 mutation back to wild-type DNA sequence, thus excluding an effect of *Mcm2* gene mutation (Fig.3G, S3-E). Interestingly, while the genes were deregulated in all analysed clones, there was a varying distribution of up-and down-regulation between clones, for example in the 4933427D06Rik, Klf15 and Iqsec1 genes (Fig. 3G), indicating stochastic de-regulation of transcription not restricted to repression. The 3D genome organization of the locus in WT cells (Fig. 3F, S3D) revealed that transcriptional deregulation spread upstream of the *Mcm2* gene to A compartment loci and was not limited by an intersecting B-compartment between the Chchd6 and Zxdc genes (Fig. 3F-G, S3D-E). Like the experiments in the c-MYC locus, we observed a distance-dependent amplitude of the transcriptional deregulation (Fig. 3G). Notably, these clones were propagated long term, so the observed effects clearly persisted through many cell divisions without recovering the wildtype expression. Collectively, the data presented so far support a model by which a single DNA double-strand break stably alters the transcriptional output from the topologically defined chromatin domain surrounding the break site and that such changes are propagated across multiple generations of dividing cells.

### Heritable alteration of c-MYC expression is accompanied by changes in the 3-D chromatin makeup

Due to the established dependence of the c-MYC locus on genome architecture (*35*, *36*) and the general notion that genomic loci undergo topological changes after DNA breakage (*24*), we set out to assess the 3-D makeup of the c-MYC locus. We reasoned that topological changes before and after DNA damage could contribute to impaired transcription at post-repair chromatin. To account for the cell-to-cell variations in 3-D chromatin arrangement (*43*– *45*) we were looking for conditions that would allow us to address this at a single-cell, or even better, a single-allele level. To this end, we employed dual-color 3-D DNA fluorescence *in situ* hybridization staining (DNA FISH), an established approach to quantify distance between genomic regions (*46*, *47*), combined with immunofluorescence labeling of the repair protein 53BP1 as a marker of the Cas9-generated DSBs in the c-MYC ORF (Fig. 4A, B). The two FISH probes were located near the boundary regions of the c-MYC TAD, covering 400 kb (Fig. 4A; upstream probe, magenta) and 450 kb (Fig. 4A; downstream probe, cyan), respectively. Consistent with the hyper-triploid nature of the HeLa cell line, these probes yielded on average four dual-colored foci per cell using 3-D confocal imaging (Fig. 4A, B). The first parameter we quantified was the distance between the magenta and cyan spots as proxy for the folding of the c-MYC TAD (Fig. 4C, D). In naïve cells, the average spot distance was at 0.48 µm, which increased during 48h after the Cas9 cut in the c-MYC ORF to 0.63 µm (Fig. 4D, S4B). This amounts to an almost 30% increase in 3-D distance between the two ends of the c-MYC TAD in cells that recovered from a Cas9-generated DSB. We next quantified the volume of the spots to approximate chromatin compaction at each single c-MYC allele (Fig. 4E). Following different kinetics after DSB generation, both regions were found in a hyper-compacted state within the 48h time-course of the experiment (Fig. 4F, S4B). Combined with the spot distance measurements, these results indicate local chromatin compaction accompanied by partial loss of the long-range interactions within the TAD as a lasting consequence of DNA breakage.

**Fig. 4.**
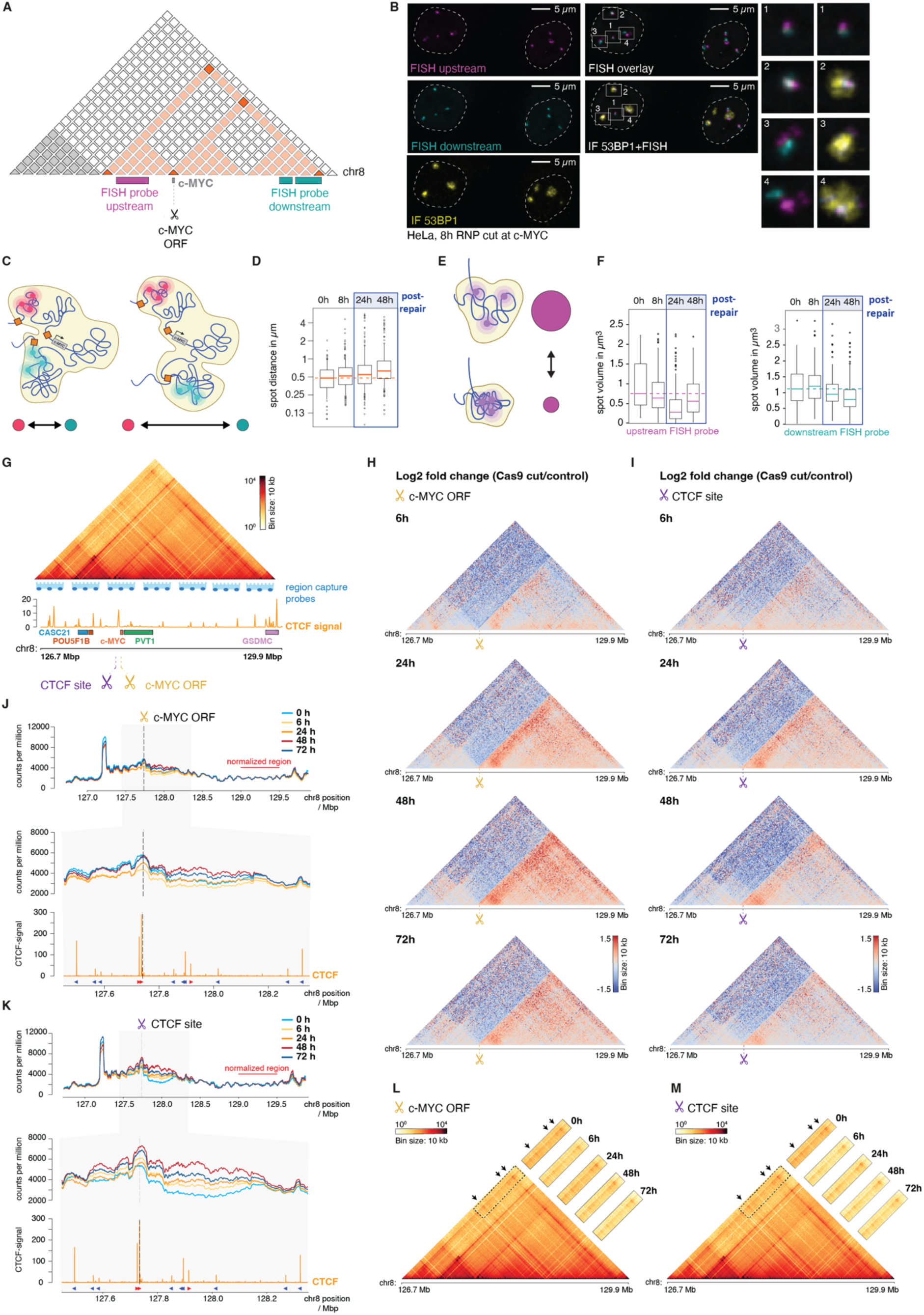
Heritable alteration of c-MYC expression is accompanied by changes in the 3D makeup of post-repair chromatin. (A) Schematic depiction of the c-MYC TAD on chromosome 8 (highlighted in orange) with indicated Cas9 cutsites (scissor symbols) and the location of the upstream (magenta) and downstream (cyan) DNA FISH probes. (B) Representative images of HeLa cells 8 hours after Cas9 RNP transfection. The respective FISH probes are pseudocolored as in (A), immunostained 53BP1 is depicted in yellow. Scalebar represents 5 µm, boxes 1-4 show magnified insets of the two FISH signals (left) with additional overlay by the 53BP1 signal (right). (C) Schematic representation of DNA FISH spot distance analysis as a measure of folding changes in the c-MYC TAD. (D) Quantification of FISH spot distances at indicated timepoints after Cas9 cut in c-MYC ORF (See also figure S4A). Plots show individual spot distances of n=102-216 spots per sample in µm of a representative experiment of two biological replicates, red bars indicate the median and the red dotted line marks the 0h timepoint. (E) Schematic representation of DNA FISH spot volume analysis as a measure of chromatin compaction changes in the c-MYC TAD. (F) Quantification of FISH spot volumes after indicated timepoints upon Cas9 cut at c-MYC ORF (see also figure S4B). Plots show individual spot volumes in µm_3_ of n=102-216 spots per sample for the upstream and downstream FISH probe, respectively. The data depict a representative experiment of two biological replicates, red bars indicate the median spot volume, the red dotted lines mark the 0h timepoint. (G) Region Capture Micro-C (RCMC) contact matrix at 10 kb resolution (top panel), and CTCF ChIP-seq profile (bottom panel). Tiling of RCMC probes within the c-Myc TAD indicated by blue probe symbols. c-Myc and CTCF cut sites are indicated by scissor symbols. (H) Fold change contact matrixes of respective time points after Cas9 cleavage at the c-Myc locus over control. All contact maps are binned at 10 kb resolution. Color scale is set to depict the loss of interactions in blue and the gain of interactions in red. Data plotted are the average of two biological replicates (see also figure S4C). (I) same as (H), but for CTCF cutsite. (J) Top panel: 1D MNase-seq profiles from Micro-C data after cleavage at the c-Myc site (indicated by dotted line). The reads within the c-Myc TAD were normalized to reads from 129.0 Mbp – 129.5 Mbp region (indicated by red bar). Middle panel: Zoom into CTCF-flanked region enclosing the cut site. Bottom panel: CTCF ChIP-seq profile at respective region. Red and blue arrows indicate CTCF motif orientation. (K) Same as (J), but for CTCF cutsite. (L) large panel: c-MYC TAD interaction heatmap at 10 kb resolution of the control experiment. Dashed rectangular indicates the position of snippets displayed in smaller panels. Snippets of trans-cut site interactions for respective timepoints after Cas9 cleavage at the c-Myc cut site. (M) Same as (L), but for CTCF cutsite.

To corroborate these findings on a cell-population base and gain high-resolution insight in the topological changes within the c-MYC TAD before and after DNA repair, we measured the dynamics of structural rearrangements with Region Capture Micro-C (RCMC)(*48*)(Fig. 4G). Over a timecourse from 6 to 72h, we observed a decrease in interaction across the respective Cas9 cutsite compared to the 0h control (Fig. 4H-I, S4C), in accordance with the dissociation of FISH foci (Fig. 4D). In addition, a progressive compaction downstream of the cut site was observed, reaching a peak intensity between 24 and 48 h after damage induction (Fig. 4H-I). Interestingly, the progressive compaction was restricted to downstream of the cut site, with more short range contacts the closer to the cutsite. These topological rearrangements were observed after both DSBs (c-MYC ORF, Fig. 4H and upstream CTCF site, Fig. 4I) at all time points in line with local chromatin compaction observed with DNA FISH (Fig. 4F), suggesting they persist beyond DNA break repair.

We further assessed the nature of DSB-induced chromatin compaction by extracting the MNase-seq information from the Micro-C dataset (*5*). We found that overall, the 1D nucleosome landscape remained unchanged (Fig. 4J-K). However, at DSB-proximal regions, we observed MNase accessibility changes which were more pronounced after cutting at the CTCF site than in the c-MYC ORF (Fig. 4J-K, middle panels). Intriguingly, even after 72h, cells did not recover the naïve MNase accessibility indicating lasting accessibility changes at break-proximal sites (Fig. 4J-K).

Importantly, our experiments revealed maintenance of CTCF-mediated loops across the DSB throughout the entire time course of breakage, repair, and recovery (Fig. 4L-M). Thus, despite the changes in long-range contacts, short-range contacts and MNase accessibility, the overarching TAD structure seems largely maintained, excluding catastrophic locus disruption.

### A single Cas9 DSB decreases RNA retention across the post-repair c-MYC TAD

To understand how the 3-D chromatin alterations of post-repair chromatin impair gene activity, we asked whether these long-term consequences of DNA breakage translate to heritable changes in functional properties of the c-MYC TAD. One function of an actively transcribed locus is to coordinate release and retention of nascent RNA, the latter of which has recently emerged as an important input to the formation of 3-D nuclear compartments (*49–53*). Indeed, there are at least two transcripts derived from the c-MYC TAD, the PVT1 lncRNA gene and the c-MYC gene itself, which were previously shown to focally accumulate at the location of their genomic origin using RNA FISH (*54*). We thus hypothesized that the post-repair changes in the c-MYC TAD might be coupled to impaired retention of the transcripts and the subsequent alterations of the nuclear compartments formed around them. To test this, we employed RNA FISH labeling of PVT1 transcripts (Fig. 5A-B, cyan) and c-MYC transcripts (Fig. 5A-B, magenta) using probes targeting the respective exonic regions. The corresponding nuclear foci also contained non-spliced species as they could be visualized using intronic probes as well (our unpublished observations). We quantified both the number and the density of the RNA foci, two parameters that notably do not represent the entire population of PVT1 and c-MYC transcripts and do not distinguish spliced and unspliced transcripts, but only the fractions retained at the locations of their origin.

**Fig. 5.**
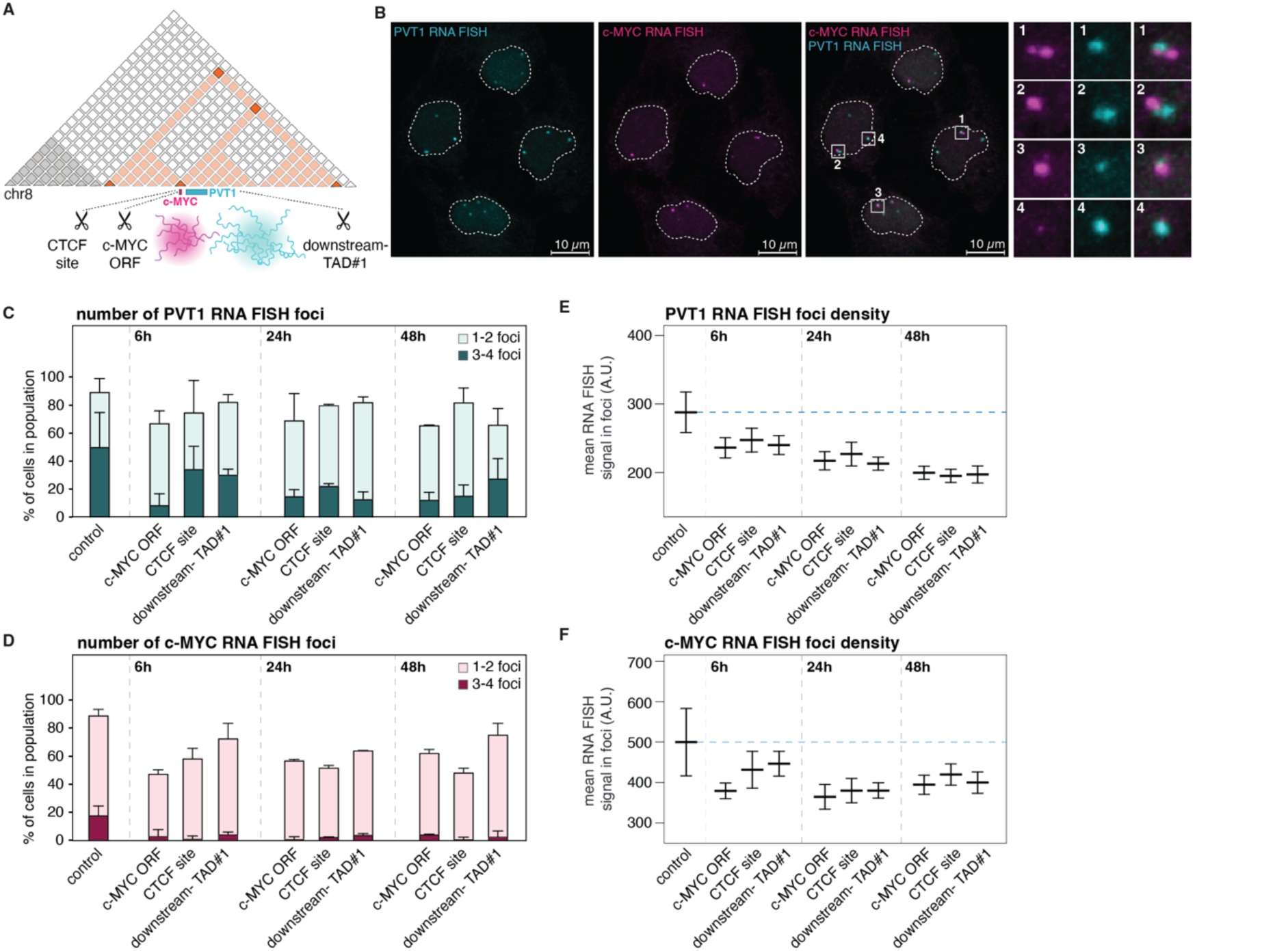
A single Cas9 DSB reduces post-repair RNA retention across the c-MYC TAD. (A) Schematic depiction of the c-MYC TAD on chromosome 8 (highlighted in orange) with indicated Cas9 cutsites (scissor symbols). The gene location and focal RNA accumulation of c-MYC (magenta) and PVT1 (cyan) are shown. (B) Representative images of RNA FISH signals in control HeLa cells without external DNA damage; the color code for the c-MYC and PVT1 RNA FISH signals is the same as in (A). Scalebar represents 10 µm, boxes 1-4 show magnifications of the individual (left, middle) and overlaid (right) signals of the FISH probes. (C, D) Quantification of PVT1 (C) and c-MYC (D) local RNA accumulation in cells with 1-2 RNA foci (light bars) and 3-4 RNA foci (dark bars) at indicated timepoints after Cas9 cutting at the indicated cutsites. Error bars are the standard deviation; n=20-70 cells per condition per replicate, data plotted is the average of two biological replicates. (E, F) Mean intensity of local RNA FISH signal of PVT1 RNA (E) and c-MYC RNA (F), horizonal bar is the mean of two biological replicates, error bars are standard deviations (see also figure S5A-B).

In naïve cells, half of the cells showed 3-4 PVT1 RNA foci and roughly 40% 1-2 PVT1 foci, while around 20% of cells had 3-4 c-MYC RNA foci and 70% 1-2 c-MYC RNA foci. Around 10% of cells had no visible PVT1-or c-MYC RNA foci (Fig. 5C, D). We observed 3-4 combined spots per cell with different amounts of each RNA species, in agreement with on average four c-MYC loci per cell we identified using DNA FISH (Fig. 4B). We then assessed whether and how these parameters change after inducing Cas9 cuts in the c-MYC ORF, the adjacent CTCF site, and one additional site downstream of both transcript origins (downstream-TAD#1; Fig. 5A). PVT1 RNA foci were partially reduced after 6h of Cas9 cut in all sites. This partial reduction of PVT1 lncRNA retention was maintained up to 48h after Cas9 cleavage (Fig. 5C). The c-MYC RNA foci became markedly reduced at all sites at the 6h time-point and did not measurably recover 24h-48h after Cas9 cutting (Fig.5D). Thus, both the PVT1 and c-MYC locally accumulated RNAs were reduced to varying degrees after Cas9 cutting. We furthermore set out to quantify the mean intensity within the RNA spots, since the density of RNA in the RNA compartments may contribute to the impaired TAD structure and function observed in our previous experiments (Fig. 1-4). Indeed, both PVT1 and c-MYC foci detectable after 6h, 24h and 48h contained a lower concentration of RNA species (Fig. 5E-F, Fig. S5A-B), consistent with lower transcriptional activity and compromised compartment formation at actively transcribed loci that have repaired a DSB in-or outside the coding sequences. Collectively, these data further support the emerging model whereby topological alterations of post-repair chromatin in the c-MYC TAD undermine full transcriptional activity and compromise RNA-based compartmentalization at sites of active transcription spanning both protein coding and non-coding RNA species. In turn, impairment of local RNA scaffold formation may explain enhanced local chromatin compaction (*55*, *56*).

### The DSB-recovered c-MYC locus remains less responsive to physiological stimulation

Finally, to investigate the physiological consequences of transcription alteration at post-damage chromatin, we asked whether a locus that recovered from DNA breakage is still proficient to properly read, and react to, physiological cues. The c-MYC gene is driven by a plethora of stimuli and as such reacts to metabolic as well as stress signaling (*57*, *58*). To test whether physiological stimulation of the c-MYC locus is still proficient in cells that have successfully repaired DSBs within the c-MYC TAD, we set out to quantify their susceptibility to respond to epidermal growth factor (EGF) stimulation, a canonical driver of c-MYC expression (*59*, *60*)(Fig. 6A, B). Specifically, we subjected c-MYC-AID-mEGFP HeLa cells to Cas9 cutting, allowed them to repair and recover for 24h, and subsequently depleted exogenous growth factors by culturing the cells in serum-free medium for another 24h (Fig. 6B).

**Fig. 6.**
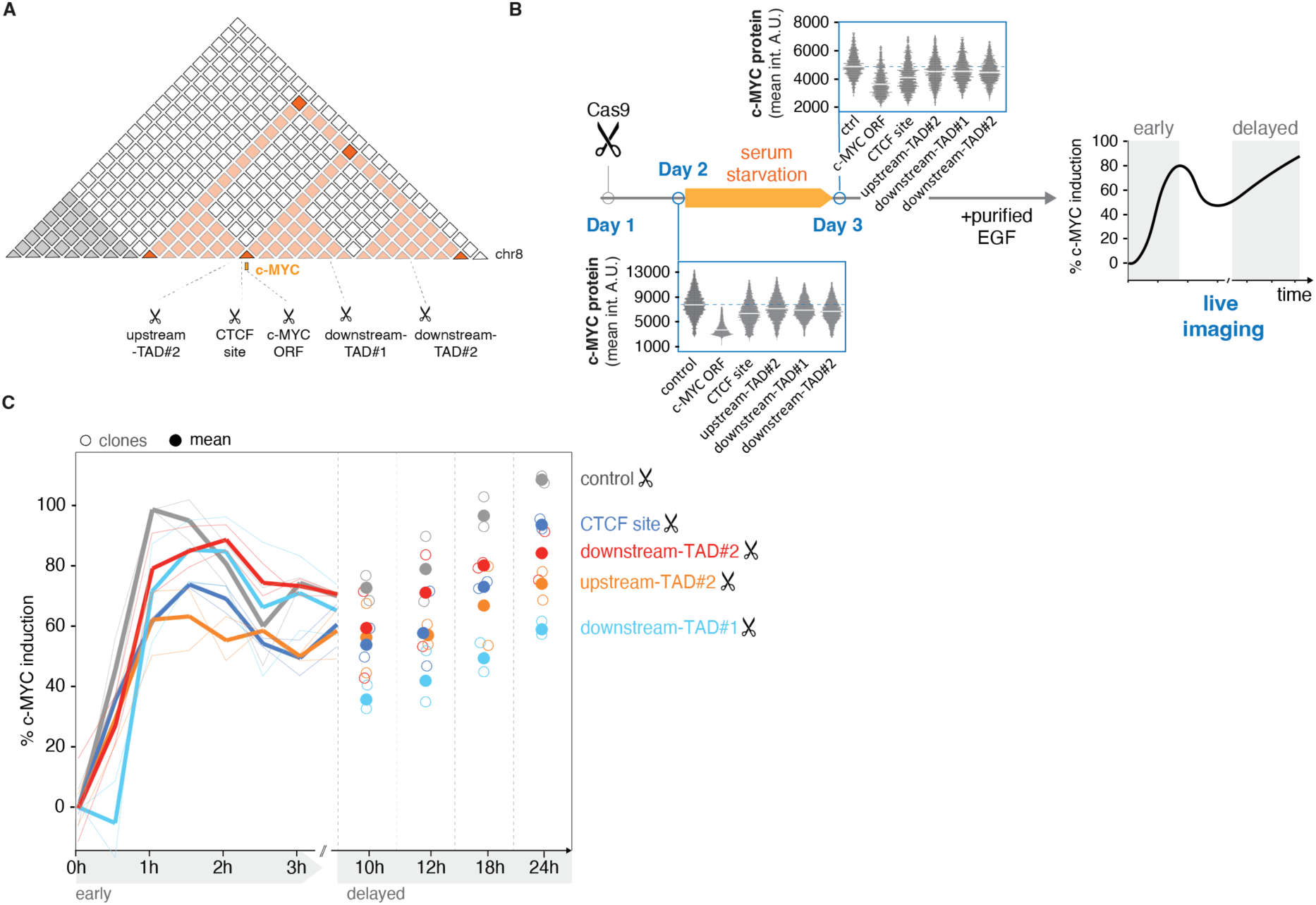
Post-repaired c-MYC locus remains less responsive to physiological stimulation. (A) Schematic depiction of the c-MYC TAD (highlighted in orange) with indicated Cas9 cutsites (scissor symbols). (B) Endogenously tagged c-MYC-AID-mEGFP HeLa cells were transfected with Cas9 RNPs, recovered for 24h, subjected to growth factor starvation in serum-free medium for 24h, and subsequently induced with 10 ng/ml purified epidermal growth factor (EGF). Immediately after EGF addition, cells were subjected to high-content live imaging for 24h. Data show mean c-MYC protein expression levels of one representative clone at the given stage of the experiment for each indicated cutsite; the dotted lines mark the control cells transfected with non-targeting Cas9. The right panel is a schematic depiction of a growth factor induction experiment highlighting early and delayed phases of c-MYC induction. (C) Relative induction of c-MYC in growth-factor depleted c-MYC-AID-mEGFP HeLa cells by purified epidermal growth factor (10 ng/ml EGF) (see also figure S6A). The scale was adjusted to the maximal induction measured in control cells, and values are shown relative to t0 (right after EGF addition). The line plot (left) shows the early boost of induction in two independent cell line clones, n=100-200 cells per clone, timepoint and condition. The dot plots (right) show the delayed gradual accumulation of c-MYC at indicated timepoints. Empty circles are different clones, full circles indicate mean.

Aligned with our previous data, the persistent reduced levels of c-MYC protein were confirmed by QIBC (Fig. 6B). We then added purified EGF and quantified c-MYC induction using high content live cell imaging. Growth factors induce c-MYC expression in two phases, an early acute burst stimulation that drops after a few hours, and a delayed gradual accumulation of c-MYC ((*59*), Fig. 6B). Indeed, we could clearly discriminate both induction phases in control cells which have not seen a DSB in the c-MYC TAD (Fig. 6C, Fig. S6A, grey). Importantly, cells which have been exposed to, and recovered from, the Cas9-generated DSBs in the c-MYC TAD consistently displayed a marked drop to 60-80% of the induction rate during the early response (Fig. 6C) as well as throughout the duration of the late response (Fig. 6C, Fig. S6A). These findings add an important functional ramification to our previous results by showing that while the extent of transcriptional impairment in genomic loci that encountered and recovered from DNA breakage may be compatible with basal cell proliferation, it severely compromises the ability of the gene regulatory elements to sense physiological stimuli and react to them accordingly. Translated to the tissue and organ context, heritable fatigue of post-repair chromatin spanning topologically defined genomic segments can undermine important decisions during development, tissue renewal in the adult organism, cell cycle commitment and others that are reliant on timely response to external or internal regulatory cues.

## Discussion

It is well-established that whenever a DSB occurs, it is in the context of 3-D folded and epigenetically defined chromatin (Fig. 7i). In addition, large segments of chromatin around the actual DNA lesion are subjected to histone modifications and 3-D restructuring (Fig. 7ii). However, it remains elusive whether chromatin functions in the affected loci recover to the pre-damaged level after restoring the integrity of primary DNA sequence. Here, we address this question and conclude that DNA repair leaves a persisting fatigue in the post-repair chromatin structure and function, which can affect the entire 3-D chromatin neighborhood (Fig. 7iii). We propose that these post-repair alterations of chromatin architecture can last throughout at least several rounds of successive cell divisions and thus prime for accumulation of potentially transforming changes in chromatin function of a cell lineage (Fig. 7iv).

**Fig. 7.**
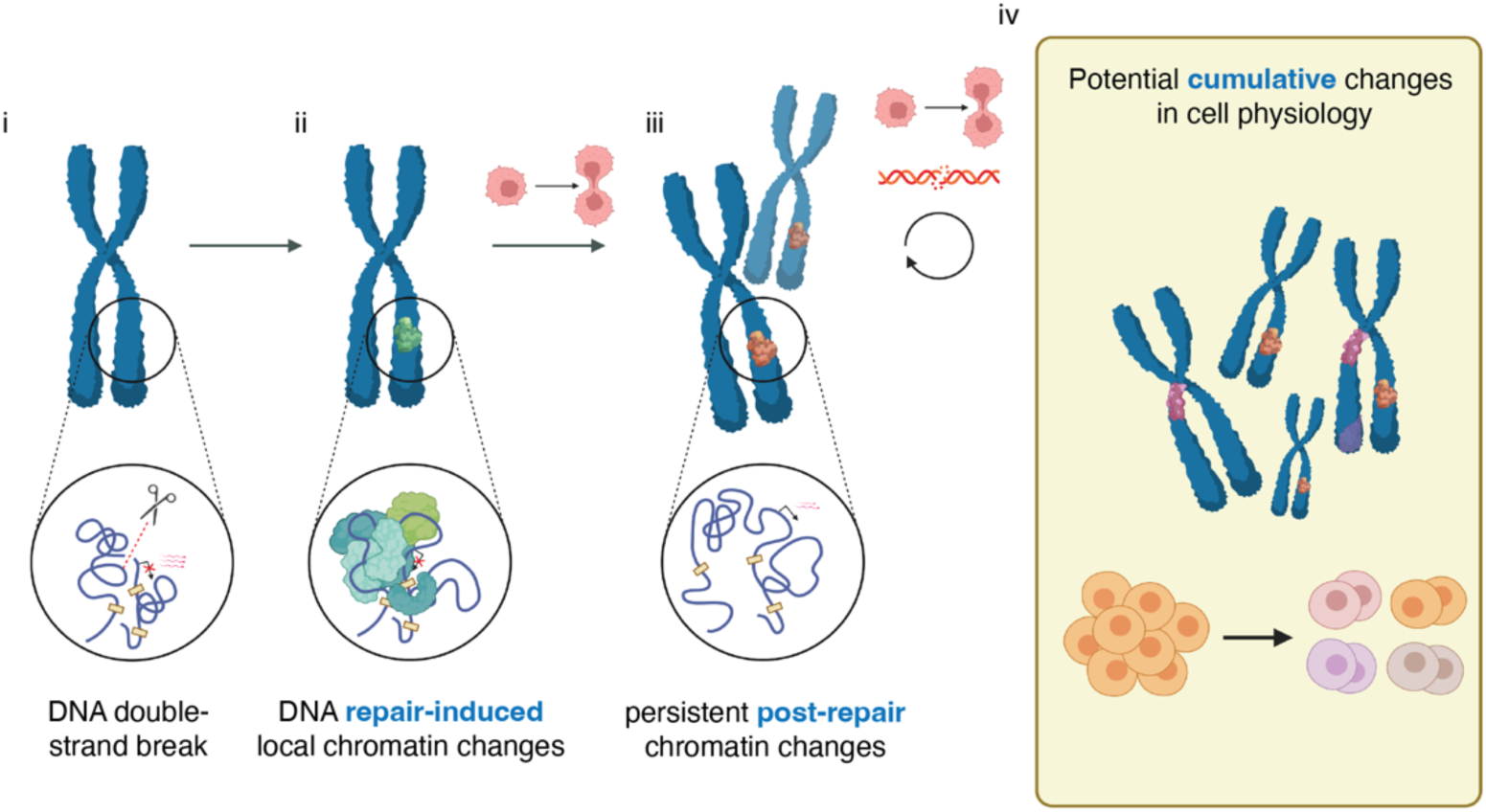
Model: Heritable consequences of DSB repair-associated chromatin alterations to cell physiology. Model figure summarizing acute and lasting changes in damaged and repaired chromatin and the potential consequences for cellular physiology. Details are in the main text.

### Post-repair chromatin fatigue as an additional layer to DNA mutagenesis

In addition to point mutations affecting single genes, our results uncover a second heritable dimension of clastogen assaults, which we term here as post-repair chromatin fatigue. This post-repair chromatin fatigue potentially affects numerous genes within the topologically defined chromatin neighbourhood that encountered, and recovered from, a single DNA breakage. This phenomenon is distinct from the recently described micronuclei-associated transcriptional repression of genes after re-integration into the nuclear genome (*61*, *62*). As reported in these studies, chromosome fragments being ejected from the nucleus in the form of micronuclei are brought into contact with cytoplasmic factors and seem to inherit persistent DNA damage accompanied by heterochromatinization and transcriptional repression (*61*). In our work, we applied Cas9 to understand consequences of single DSBs that could be fully repaired in otherwise intact interphase nuclei. In such settings, we do not detect lasting DNA damage or chromatin modifications associated with heterochromatinization (our unpublished observations) but we consistently see lasting transcriptional attenuation in the progenies of mother cells exposed to single DSBs. Based on our data, we propose that this is rooted in the lasting chromatin topology changes invoked by breakage of both DNA strands, which disrupt the natural 3-D environment and thus the physical and functional interplay between chromatin elements that determine the physiological dynamic range of gene expression. Thus, the final inheritance of deregulated transcription observed in both scenarios is shared but follows different strings of causative events.

### Potential causes of heritable alterations in post-repair chromatin

How could successful repair of a DSB lesion lead to lasting topological changes? It has been shown that chromatin around a DSB undergoes a massive 3-D restructuring converging on transcriptionally silent domains that are, at least in part, defined by adjacent TAD boundaries and whose spatial arrangement related to the neighbouring TADs is stabilized by dedicated factors including the cohesin complex, 53BP1, RIF1 and SLFN5 genome caretakers, or ASF1 histone chaperone (*24*, *28*, *30*, *31*, *63*). These previous studies showed that DNA repair process *per se* can benefit from such chromatin restructuring (e.g. by locally concentrating ultra-low abundant repair factors that foster the fidelity of DNA repair (*24*, *64*)), but our current study revealed that this may come with the ‘price-tag’ of lasting rearrangements and chromatin compaction known to hamper transcription factor binding (*65*, *66*). We envisage that such unscheduled dampening of gene activity could reinforce itself by seeding heritable alterations of chromatin topology that would preclude its return to the pre-damaged state. Indeed, intriguing conceptual analogies supporting such hypothesis have been recently reported and include the ability of IFN response genes to memorize stimulations that are also instigated by *de novo* chromatin alterations (*11*). In extension of this model it has recently been suggested that sampling frequencies between regulatory elements can be affected by altered local transcriptional activity (*67*). It is thus possible that fatigued chromatin displays different conditions for such sampling manifesting irregular transcriptional regulation.

Furthermore, DSBs can trigger changes in the chromatin landscape by untimely or excessive accrual of topology regulators. Prominent among those is CTCF, a canonical TAD insulator, whose known accumulation at DSB-flanking chromatin microdomains (*68*, *69*) is consistent with transcriptional deregulation described in our study. Firstly, CTCF anchor engagement per se seems enhanced under conditions of RNAPII removal (*70*), such as the one that can occur at damaged chromatin (*71–73*). Additionally, several non-coding RNAs have been implicated with CTCF eviction to regulate local chromatin folding (*74–76*), consistent with the possibility that reduced retention of RNA species at post-repair chromatin as shown in our study could foster ectopic CTCF accumulation. In such scenario, the hyper-loading of CTCF and subsequent stiffening of the affected locus might reconfigure spatial relationships between genes and their enhancers, which may not cause complete transcription shutdown but rather lead to self-perpetuating contraction of the transcriptional dynamic range. Our results showing the inability to respond to mitogenic stimulation and mount physiological levels of c-MYC after repair of DSBs that do not affect its coding sequence but are located in the same TAD are consistent with such reasoning. Of note, changes in local chromatin folding and compaction by the above-described mechanisms is likely prone to stochasticity whereby some promoter-enhancer pairs might be locked, while others could get disrupted. In line with this idea are our observations in the post-repair MCM2 locus, which mainly shows inherited transcriptional repression, yet with consistent upregulation of a subset of local transcripts. Additionally, intriguing conceptual analogies exist in different biological contexts such as X chromosome silencing where TAD inversion by DSB-generating nucleases revealed similar randomization of surrounding gene activity for the XIST locus (*77*).

As alluded to in the previous paragraph, an exciting aspect of post-repair chromatin uncovered by our work is reduced retention of at least two RNA species known to locally cluster at the c-MYC TAD. RNA contributes to genome structure in multiple ways, ranging from specific local rearrangements to RNA-based structural support of chromosome architecture (*78*). Perhaps most relevant to our results, local accumulation of both coding and non-coding RNA species has been reported to seed regulatory micro-compartments (*49*, *79*), which modulate the local chromatin density via various means including 3-D genome topology (*55*, *56*, *80*, *81*). It is thus tempting to speculate that the observed inability of post-repair chromatin to focally concentrate c-MYC and PVT1 RNA species may contribute to lasting changes in 3-D chromatin environment that in a heritable fashion remodel the local environment for gene activities across the affected TAD. In addition, a recent study has shown wide-spread association of RNA species with transcription factors modulating dynamics and specificity of their action (*82*). It is thus similarly possible that reduced transcription after break repair causes defects in local TF regulation.

Finally, an important mechanistic ramification of heritable fatigue of post-repair chromatin includes passage through mitosis. On the one hand, the extensive DNA compaction during mitosis and re-formation of the interphase nucleus in the subsequent G1 phase (*39*) could provide a window of opportunity for the cell to reset the chromatin state and reverse potential alterations in chromatin structure introduced during DSB repair. On the other hand, our findings raise the possibility that the *de novo* DSB-induced changes in 3-D chromatin makeup could be stabilized by mechanisms similar to mitotic bookmarking, which manifest in stabilized chromatin accessibility patterns to enforce physiological chromatin memory (*83*, *84*). Indeed, at least one key component of transcriptional mitotic bookmarking, the SWI/SNF chromatin remodeler SMARCE1 was recently shown to be among proteins that are recruited to DSB-flanking chromatin (*85*).

### Functional implications of lasting rearrangements of post-repair chromatin

Context-specificity is an important aspect of our findings. While not all genes may be regulated by topological features, it seems clear that in specific cases, such as the c-MYC locus, local topology has evolved to support the physiological expression patterns. For c-MYC, this is manifested in a 3-D enhancer network spread over the entire TAD to interact cooperatively in a cell-type-specific manner (*86*). As c-MYC is crucial for cellular proliferation but also drives tumour growth if overexpressed, it’s TAD is a canonical example of a regulatory hub where the 3-D makeup of the surrounding chromatin plays a defining role in conferring transcriptional robustness. In this context, as we show, even a single repaired DSB can cause lasting physiological perturbations. Viewed from this perspective, our findings can have broader implications for predicting context-specific consequences of DNA breakage. We envision that housekeeping genes do not heavily rely on genome topology to sustain their stable expression and thus have a low risk of interference by post-repair chromatin fatigue. In contrast, genes which acutely sense regulatory cues and propagate signaling pathways during metabolic fluctuations, stress stimulation, differentiation, evolution, or cell cycle transition are much more likely to leverage the plasticity of the surrounding 3-D genome context to regulate their expression. Hence, these genes will have a higher likelihood to be susceptible to interference by damage-induced chromatin mutagenesis. Due to the stochastic nature of the DSB-induced topological rearrangements the extent of transcription deregulation does not need to be instantly high, but when accumulating over the genome during the lifetime of an organism, it could lead to progressive decline of genome function and contribute to tissue ageing. One such example could be multistage carcinogenesis, where a single DSB can derail cellular evolution in two ways. In parallel to the progressive accumulation of DNA sequence mutations in tumour suppressor genes, we speculate that in at least some parts of the genome, the topological alterations of post-repair chromatin likewise permanently modify responsiveness of those tumour suppressors, whose genes are located within the topologically defined chromatin neighbourhood that encountered, and recovered from, DNA breakage, which are hallmarks of unstable genomes and whose incidence progressively increases during cancer progression.

Finally, the concept of post-repair chromatin fatigue has implications for gene editing using DNA double-strand breaks to modify the genome, most notably the CRISPR-Cas9-mediated approaches. Our findings of heritable dysregulation of c-MYC dynamic range after Cas9-mediated DSB anywhere in the c-MYC TAD as well as altered transcriptional output in an entire genomic region around endogenously modified MCM2 in mouse embryonic stem cells indicate that caution should be exercised when considering the physiological outcomes of genome editing. Our results suggest that it is worth to inspect whether the Cas9-target gene resides in topologically sensitive chromatin neighbourhood, which other genes are located nearby, and verify that phenotypes strictly derive from the intended mutation and not from alterations of adjacent gene(s) due to post-repair chromatin fatigue.

## Acknowledgements

Research funding was provided by the Novo Nordisk Foundation (grant NNF14CC0001). Irene Mordini and Nils Krietenstein were supported by the Lundbeck Foundation (grant R368-2021-1076) and the Novo Nordisk Foundation (grant NNF14CC0001). Research in the Groth lab is supported by the European Research Council (ERC CoG 724436), the Lundbeck Foundation (R198-2015-269 and R313-2019-448), Independent Research Fund Denmark (7016-00042B; 4092-00404B), and the Novo Nordisk Foundation (NNF21OC0067425). Alva Biran was supported by Marie Curie Individual Fellowships (846375). We thank Maj-Britt Rask for expert technical assistance and Natalia Frese for lab maintenance. We thank the Protein Imaging Platform at the Novo Nordisk Foundation Center for Protein research for support with microscopy and image analysis. We thank the Flow Cytometry Platform at the Novo Nordisk Foundation ReNew Center for FACS sorting, and the CPR/reNEW Genomics Platform for RNA sequencing and Region Capture Micro-C sequencing. We thank all members of the Lukas laboratory for conceptual and technical inputs to this study and for discussions and critical comments on the manuscript. Model figures were generated using BioRender.com.

## Supplemental information

### Materials and methods

#### Cell culture

Cells of the human HeLa Kyoto cervical cancer cell line (obtained from S. Narumiya) were grown under standard cell culture conditions (5% CO_2_, humidified atmosphere) in Dulbecco’s modified Eagle’s medium (DMEM)(ATCC, 30-2002) containing 10% heat-inactivated FBS and penicillin–streptomycin antibiotics (Thermo Fisher Scientific, Invitrogen 15140-122). The following cell lines with genetic modifications were used: HeLa cells endogenously tagged cell lines c-MYC-EGFP-mAID clones 23 and 32 (Fig. S1G, Fig. 6B-C, Fig. S6A). Cells were regularly tested for absence of mycoplasma (MycoAlert, Lonza, BioNordika LT07) and authenticated by STR profiling (IdentiCell Molecular Diagnostics). For epidermal growth factor (EGF) stimulation of cells, 10 ng/ml purified EGF (Gibco, ThermoFisher PHG0311L) was added to cells after 24h growth factor depletion in serum-free medium (Fig. 6 and Fig. S6).

#### Generation of DNA double-strand breaks using Cas9 RNPs

To generate site-specific DNA double-strand breaks, cells were transfected with sgRNA–Cas9 ribonucleoprotein complexes (RNPs) using Lipofectamine CRISPRMAX Cas9 (Invitrogen, CMAX00008). All guide RNAs used were single guides (IDT) and assembled to RNPs with TrueCut Cas9 Protein V2 (Invitrogen, A36499), nickase Cas9D10A (IDT, 1081062) or nickase H840A (IDT, 1081064) according to manufacturer’s instructions. For transfection of a 60-mm dish (3 ml), 0.9 μl Cas9 enzyme (5 mg/mL stock) was diluted in 136 μl Opti-MEM medium followed by addition of 9 μl sgRNA (2 μM) and 18 μl Plus-Reagent from the CRISPRMAX kit. CRISPRMAX reagent (12 μl) was separately diluted in 170 μl Opti-MEM medium, subsequently added to the RNP reconstitution, incubated for 5-10 minutes at room temperature and added to cells.

#### Antibodies

Antibodies against the following antigens were used: anti-53BP1 (mouse, Millipore, MAB3802, 1:200 for IF), anti-c-MYC (abcam, ab32072, 1:2000 for IF, 1:1000 for WB), anti-GFP (rabbit, Proteintech group, PABG1, 1:3000 for IF, 1:2000 for WB). All antibodies were validated by the manufacturer. Secondary antibody conjugates for fluorescent detenction were goat-anti mouse and goat-anti rabbit Alexa Fluor 488 (Invitrogen, Thermo Fisher A11029; A11034), Alexa Fluor 568 (Invitrogen, Thermo Fisher A11031; A11036), Alexa Fluor 647 (Invitrogen, Thermo Fisher A21236; A21245) reagents (Thermo Fisher Scientific).

#### Immunofluorescence (IF)

Cells were grown on cleaned and autoclaved glass coverslips to a confluency of around 60% for Cas9 RNP transfections, and treated as indicated. Treated cells were fixed in 4% formaldehyde for 12 min at room temperature, and subsequently permeabilized for 5 min in PBS and 0.2% Triton X-100 (Sigma-Aldrich, T9284). All antibodies were applied in filtered DMEM (ATCC, 30-2002) containing 10% FBS. Primary-antibodies were incubated at room temperature for 2 hours. Coverslips were then washed with 1xPBS (FisherScientific, Gibco, 14190-169) containing 0.2% Tween (Sigma-Aldrich, P2287). Secondary antibodies were incubated at room temperature for 1 hour and were supplemented with 4ʹ,6-diamidino-2-phenylindole dihydrochloride (DAPI, Molecular Probes, D1306, 0.5 μg/ml) to label DNA. After three more washes in PBS-Tween, coverslips were washed twice in distilled water, air-dried for at least one hour, and mounted in 6 μl Mowiol-based mounting medium containing Mowiol 488 (MilliporeSigma Calbiochem, 475904100GM).

#### QIBC

Image acquisition for QIBC by high-content Widefield microscopy (ScanR inverted high-content Screening station, Olympus) was performed as previously described (*87*). Images were captured using widefield optics, a UPLXAPO dry objective (20×0.80-WD0.60mm), Lumencor Spectra LED light source, SPX-QSEM Quadband Fluorescence mirror unit and dichroic beamsplitter, and a sCMOS camera (Hamamatsu ORCA flash4) chip. For each sample, at least 1000 but usually >2000 cells were imaged and analysed with the ScanR analysis software (Olympus Corporation, ScanR analysis software versions 2.8 and 3.2). Number and intensities of nuclei were quantified with single and calculated parameters. These values were then exported for visualization using TIBCO Spotfire desktop software (TIBCO software Inc., version 12.1.1).

#### T7E1 surveyor assay

For confirmation of specific Cas9 cutting, 1 mio cells transfected with sgRNA-Cas9 RNPs for 24h were harvested in ice-cold PBS, and genomic DNA was extracted using the DNeasy Blood and Tissue kit (Qiagen, 69506) according to the manufacturer’s instructions. The target region was amplified by PCR using the CloneAmp HIFI PCR mix (Clontech, BioNordika 636298) according to the instructions. Oligos for target site amplification were designed with PrimerBLAST (NCBI, (*88*))(c-MYC ORF 1 and 2 fwd: AGCAGCCTCCCGCGACGATGC, c-MYC ORF 1 and 2 rev: GCCAATGAAAATGGGAAAGG; CTCF site fwd: GCTAAAGCTTGTTTGGCCGT, CTCF site rev: ACTTAGCTAGTTGCCCAGCC, promoter-CTCF site fwd: GACTCTTGATCAAAGCGCGG, promoter-CTCF site rev: GGGCAAAGTTTCGTGGATGC, upstream TAD#1 fwd: CCCATAACAGACATGGAATACGG, upstream TAD#1 rev: GGAACACCCCACTTCTTGGT, upstream TAD#2 fwd: CGCTTCCAAGCAGGTTTGTT, upstream TAD#2 rev: GGTCATCTTTCACCCTGCGA, downstream TAD #1 fwd: GAGCCTCCCTGAAGGCTTTT, downstream TAD#1 rev: TGGCCAACATGGTGAAACCT, downstream TAD #2 fwd: AACAAAGAGGCATTCAAGTTGCT, downstream TAD#2 rev: TTCCCCCGACGTAAACAAAG, PVT1 fwd: TTGCATACTGGCAGCGACAA, PVT1 rev: GCCAGAGAAACGTGTCCATC, FAM84B fwd: TTGCGTCGCTTCTCCATGAT, FAM84B rev: CTCGCTACACCCGAGACTTC, FAM49B fwd: ATCCTCAGGCCCTTGGTTAGT, FAM49B rev: TCCTATGGAGTGTCTGTGGAGT, POU5F1B fwd: TGGCATTCTTATCCACAAAGTGAA, POU5F1B rev: AGCCCAGAGTGATGACGGA, GSDMC fwd: CCGGGGTTTGGTATTAGTGG, GSDMC rev: CACGTTCGGTGTGAATACCC. The PCR products were run on a 1% agarose gel (Invitrogen, LifeTechnologies 16500-500), purified using the QIAquick gel extraction kit (Qiagen, 28706) according to the manual. 400 ng PCR product were subjected to a denaturing and reannealing cycle (5 minutes, 98°C with a ramp of 2°C /minute, 10 min, 25°C) in 1x NEB buffer2 (from T7E1 enzyme, NEB, BioNordika, M0302L). The DNA was split in two, half was loaded on a 1.5% agarose gel as input sample, the other half was digested with 1 µl T7E1 endonuclease (T7E1 enzyme, NEB, BioNordika, M0302L) for 30 minutes at 37°C and analysed by agarose gel electrophoresis. Gel source data is deposited.

#### Live analysis of c-MYC expression

To assess the transcriptional response to Cas9 cutting within the c-MYC TAD (Fig. S1G) and the c-MYC induction by epidermal growth factor in serum-depleted cells (Fig.6 and Fig. S6), endogenously tagged HeLa c-MYC-AID-mEGFP cells were imaged live. Cells were grown and imaged in a Cell Carrier Ultra plate (Perkin Elmer, 6057302). For nuclear tracking and segmentation, DNA was labeled with the SPY-650 DNA dye (Spirochrome, tebu bio sc-501). Widefield images were acquired using the Perkin Elmer Opera Phenix screening microscope equipped with two large-format sCMOS cameras and an OEM Andor Zyla camera (custom modified Zyla 5.5),a 20x water/NA1.0 plan Apochromat objective and temperature control and CO_2_ control modules. Laser light sources for green (488 nm) and far red (640 nm) were used. Images were acquired at indicated timepoints using the Harmony software (version 4.9). For analysis, nuclei were segmented using the Harmony software (version 4.9) and exported values of nuclei counts and mean signal intensities per nucleus were plotted in TIBCO Spotfire desktop software (TIBCO software Inc., version 12.1.1).

#### Generation of endogenously tagged HeLa c-MYC-EGFP-mAID cell lines

HeLa cells expressing C-terminally endogenously tagged c-MYC were generated using CRISPR– Cas9 as previously described (*89*, *90*). Paired single guide RNAs (sgRNAs) for the c-MYC locus (guide 1, CATCCTGTCCGTCCAAGCAG; guide 2, TGTATGCTGTGGCTTTTTTA) were inserted into pX335-U6-Chimeric_BB-CBh-hSpCas9n(D10A) (Addgene plasmid 42335, a gift from F. Zhang) using BbsI restriction. The donor plasmid carrying the tag (AID–mEGFP), a flexible linker, and flanking homology regions was synthesized as long standard gene in the pEX-A258 backbone from eurofins genomics. The plasmids carrying the nickase and sgRNA sequences as well as the homologous donor plasmid were co-transfected using Lipofectamine LTX Plus reagent (Thermo Fisher Scientific, 15338-100). GFP-positive cells were enriched by FACS. After 5 days, sorted cells were seeded into 10 cm dishes and allowed to grow single colonies, which were picked and cultured for characterization by Western blotting and junction PCR. Functional validation of positive clones was performed by immunofluorescence and Western blotting (sub-cellular localization and expression levels).

#### Western blotting

Protein analysis by western blotting was performed using standard procedures and ECL-based chemiluminescence detection. Around 1 million cells were collected from 6 well plates and lysed in RIPA buffer (50 mM Tris-HCL, pH 8.0, 150 mM NaCl, 1.0% IGEPAL CA-630, 0.1% SDS, and 0.1% Na-deoxycholic acid, protease inhibitors (Roche, 04693116001), phosphatase inhibitors (Roche, 04906837001) and 750 units per mL benzonase (Sigma, E1014-25KU)). These whole cell extract were cleared by centrifugation and analyzed by SDS–PAGE using standard procedures. Primary antibody incubation was done overnight at 4°C in blotting buffer (1xPBS (FisherScientific, Gibco, 14190-169), 0.2% Tween (Sigma-Aldrich, P2287), 5% powdered milk (Fluka, Sigma Aldrich 70166)). For primary antibody detection by chemiluminescence, secondary peroxidase-coupled antibodies (Vector laboratories, VWR PI-1000, and PI-2000) were incubated at room temperature for 1 hour, washed and subsequently immersed in ECL (Amersham, RPN2106) for analysis with an Odyssee-Fc system. Gel source data is deposited.

#### Cell cycle analysis by EdU labeling

For Click-iT EdU labeling, treated cells were incubated in medium supplemented with 20 μM EdU (5-ethynyl-2’-deoxyuridine)(ThermoFisher, 10044) for 15 minutes and then washed in 1xPBS and fixed with 4% formaldehyde (12 minutes, room temperature). Cells were first incubated with primary antibody for 2 hours, washed, and subsequently subjected to the EdU Click-iT reaction using the Click-iT EdU Alexa Fluor 647 Imaging Kit (Molecular Probes,

ThermoFisher C10340) according to the manufacturer’s recommendations. Cells were then incubated for 1 hour with secondary antibody supplemented with 4ʹ,6-diamidino-2-phenylindole dihydrochloride (DAPI, Molecular Probes, D1306, 0.5 μg/ml) and imaged by QIBC.

#### Reverse Transcription and Quantitative PCR

Total RNA was isolated using RNeasy Mini Kit according to the manufacturer’s instructions (Qiagen, 74104). The cDNA was then synthesized from 1-5 µg total RNA using the Maxima First Strand cDNA Synthesis kit (Thermo Fisher, K1641). Transcripts were quantified by real-time quantitative PCR (qPCR) using exon-spanning primers designed with PrimerBlast (*88*) (GAPDH fwd: AGCCACATCGCTCAGACAC, GAPDH rev: GCCCAATACGACCAAATCC, c-MYC fwd: GGGAGGCTATTCTGCCCATT, c-MYC rev: TAACGTTGAGGGGCATCGTC, PVT1 fwd: CTTGCGGAAAGGATGTTGGC, PVT1 rev: GCCATCTTGAGGGGCATCTT, CASC21 fwd: CCAAGAGAAGACGTCCAGCA, CASC21 rev: AGGCCAACAGGAACCACATC, POU5F1B fwd: AAGACCATCTGCCGCTTTGA, POU5F1B rev: ATCTGCAGTGTGGGTTTCGG, FAM84B fwd: GCCGAGCCTACACCTTCAAA, FAM84B rev: GGACAGGGGCTGAGGCTA) and Power SYBR Green Mastermix (Applied Biosystems, Fisher Scientific 4367659). RNA enrichment was analysed using the ΛλΛλCt method relative to the transcript levels of GAPDH and normalized to the control as indicated.

#### siRNA depletion of c-MYC

Transfection of c-MYC siRNA (Ambion Silencer Select, Invitrogen, #s9129) was performed using Lipofectamine RNAiMAX (Thermo Fisher Scientific, 13778075) at a concentration of 10 nM. Ambion negative control #9 was used as control siRNA. c-MYC siRNA depletion was performed for 24 hours, and efficient depletion was confirmed using RT-qPCR on the RNA level (primers were c-MYC fwd: GGGAGGCTATTCTGCCCATT, c-MYC rev: TAACGTTGAGGGGCATCGTC) and using QIBC on the protein level.

#### RNA-sequencing of MCM2-edited mouse embryonic stem cells (mESCs)

We re-analyzed the publicly available RNA-seq data (GSE154390) containing WT, 8 MCM2-2A (#1-#8) independent, genome edited clones, and 1 MCM2-R#2 clone. Data analyses was performed using R (v. 4.3). Differential expression analysis between WT, MCM2-2A and MCM2-R clones was performed with Deseq2 (v. 1.30.1) accounting for batch effects detected in PCA analysis, product of different sequencing runs. Differentially expressed (DE) genes were defined as those with |FC| > 1.5 and FDR < 0.01 and genes that were not rescued are defined as genes which are DE in both MCM2-2A vs. WT and MCM2-R vs. WT. Dot-plots were plotted using ggplot R package (v. 3.4.1). Mouse embryonic stem cells Micro-C based compartment analysis (4DNFILGE5LQU; (*41*)) was retrieved from the 4DN Data portal and visualized using the HiGlass genome browser (*91*).

#### DNA FISH analysis of c-MYC TAD topological changes

For DNA FISH detection of the two distal regions of the c-MYC TAD, DNA FISH was performed according to the manufacturer’s instructions (CytoCell) with minor variations. Cells were grown on coverslips, washed with PBS, and fixed in freshly made ice cold 3:1 methanol: acetic acid (methanol: Sigma Aldrich, 322415; acetic acid: Sigma Aldrich, ARK-2183) for 20 minutes at room temperature. To dehydrate cells in an ethanol dilution series, coverslips were immersed in 70% ethanol for 2 minutes, in 85% ethanol for 2 minutes and in 100% ethanol (Merck, VWR 1.11727.1000) for 2 minutes, all incubations at room temperature. Coverslips were then air-dryed for 10 minutes and pre-warmed for 10 minutes at 37°C together with the DNA FISH probes (CytoCell, LPS027). Probes and cells were then co-denatured at 80°C for 5 minutes and allowed to hybridize in a wet chamber at 37°C overnight. Finally, cells were washed once in 0.4xSSC (pH7.0)(Thermo Fisher, 15557044) at 72°C for 2 min, in 2xSSC, 0.05% Tween-20 (Sigma Aldrich, P2287) at room temperature (pH7.0) for 30 seconds and subjected to immunofluorescence staining or mounted directly after 5 minutes of incubation with PBS-DAPI (Molecular Probes, D1306, 0.5 μg/ml). For mounting of FISH-labeled coverslips, 6 µl solid mounting medium (Abberior, MM-2013-2X15ML) was used per coverslip and allowed to harden overnight. Cells were then imaged in z-stacks using the LSM880 confocal microscope (Zeiss LSM880) and images were analysed using the Imaris Cell Analysis software (version 10.0.0).

#### Confocal Imaging

For 3-dimensional confocal imaging of DNA FISH and RNA FISH samples, a confocal laser scanning microscope (Zeiss LSM880 system) equipped with a Plan-Apochromat 40×1.3 oil DIC UV-IR M27 objective, two conventional PMTs and one 32PMT GaAsP detector for fast spectral scanning, and laser lines 405, 488, 561 and 640nm were used. Z-stacks of 10-18 planes at 0.5 µm intervalls were acquired sequentially. Filter sets used were GFP (ex 470/40, em 525/50), far red (ex633, em641-695) and red (ex 550/25 em 605//70). All images were acquired by the acquisition software ZEN BLACK (Zeiss). Images were converted for analysis using the Imaris File Converter (version 10.0.0) and analysed using the Imaris Cell Analysis software (version 10.0.0). In short, DNA FISH spot foci center were segmented by automated detection and the shortest distance between two FISH spots was calculated and plotted using TIBCO Spotfire desktop software (TIBCO software Inc., version 12.1.1). Volumes of foci were segmented using an intensity mask based on absolute intensity and background correction, and the mean signal intensity within these volume masks was derived and plotted using the TIBCO Spotfire desktop software (TIBCO software Inc., version 12.1.1). For RNA FISH analysis, spot foci were segmented and counted for analysis of foci numbers per cell. RNA FISH data was plotted using Prism (Graphpad, version 9.4.1).

#### Region Capture Micro-C (RCMC)

The protocol was adapted from Goel et al. (2023)(*48*) with minor variations. Data presented come from merging two biological replicates per studied condition. For each biological replicate, 10 mio crosslinked cells were used to build the Micro-C library and successively 4 μg of Micro-C library were used to capture the locus of interest.

#### Crosslinking

For the double crosslinking, harvested cells were resuspended in 1X DPBS reaching a final concentration of 1 mio cells per mL, and crosslinked with 1% formaldehyde for 10 min at RT. The reaction was quenched with 0.25 M of glycine (final concentration) for 5 min at RT. Cells were then washed with 0.1% BSA (NEB, #B9000S) in 1X DPBS and resuspended in the same buffer at a final concentration of 4 mio cells per mL. For the second crosslinking medium a 0.3 M stock solution of ethylene glycol bis(succinimidyl succinate) (EGS) (ThermoFisher, #21565) and Dimethyl Sulfoxide (DMSO, Sigma Aldrich,#D8418) was prepared. Cells were incubated with 3 mM stock solution (final concentration) for 40 min at RT and then incubated for 5 min at RT with glycine at a final concentration of 0.4 M. After centrifugation, cells were resuspended in 0.1% BSA (NEB, #B9000S) in 1X DPBS at a final concentration of 5 mio cells for mL, divided into 5 mio and 1 mio aliquots, centrifuged, snap-frozen in liquid nitrogen, and stored at −80° C.

#### MNase titration

Different enzyme concentrations were tested to obtain the ideal DNA fragment distribution after MNase digestion. For every time point of the two biological replicates, 1 mio aliquot of frozen cells was resuspended in 500 μl 0.1% BSA (NEB, #B9000S) in 1X DPBS and incubated on ice for 20 min. Cells were collected by centrifugation at 1000 x g for 5 min RT and pellets were washed once with 500 μl MB#1 (10 mM Tris-HCl, pH 7.4, 50 mM NaCl, 5 mM MgCl_2_, 1 mM CaCl_2_, 0.2% 0.2% IGEPAL CA-630 (Sigma Aldrich, #18896), 1x Halt Proteinase inhibitor (ThermoFisher, #78430)). Cells were resuspended in 200 μl MB#1 and split into four tubes. Chromatin was digested at 37° C for 10 min 800 rpm shaking. MNase (Worthington, #LS004798) was diluted in 10 mM Tris-HCl, pH 7.4, and four different enzyme concentrations ranging from 20-0.314 U were tested. The reaction was stopped by adding 200 μl of freshly prepared STOP buffer composed of 150 μl Tris-HCl pH 7.5, 25 μl 10% SDS (Invitrogen #15553-035), 25 μl 20 mg/ml Proteinase K (GoldBio, #P-480-1), 2 μl EGTA (BioWorld, #40121266-1) and samples were incubated at 65° C for 2 hours shaking 800 rpm. DNA was extracted using Phenol-Chloroform-Isoamyl Alcohol (PCI, Invitrogen, #15593-031), purified with a commercial PCR purification kit following the manufacturer’s instructions (QIAquick PCR purification kit, QIAgen, #28106) and eluted in 15 μl of elution buffer. To define the correct MNase concentration, digested chromatin was loaded on 1.5% agarose gel and ran at 120 V for 1 hour.

#### Micro-C XL library

Micro-C XL protocol was adapted from Hsieh et al., 2020 (*41*). To decrease the duplication rate, 10 mio cells per timepoint were used to build the Micro-C library. The cells were thawed on ice for 10 min, resuspended in 1 ml 0.1% BSA (NEB, #B9000S) in 1X DPBS, and incubated for 20 min on ice. BSA was added to avoid cell clumps and reduce the cell stickiness to the tub walls. Cells were then centrifuged at 1000x g for 5 min at RT. Pellets were washed in 500 μl of MB#1 (10 mM Tris-HCl, pH 7.4, 50 mM NaCl, 5 mM MgCl_2_, 1 mM CaCl_2_, 0.2% IGEPAL CA-630 (Sigma Aldrich, #18896), 1x Halt Proteinase inhibitor (ThermoFisher, #78430)) and centrifuged at 1000x g for 5 min at RT. Cells were then resuspended in 1 ml of MB#1 and split into 1 mio cells per aliquot. Chromatin was digested with MNase for 10 min at 37° C with shaking. The proper MNase concentration was previously established via MNase titration. During this step, crosslinked frozen cells were digested with different enzyme concentrations ranging from 20-0.314 U. The condition where there are mainly mono-nucleosomal chromatin fragments was selected for future digestion.

The enzymatic reaction was stopped with 1.6 μl 0.5 M EGTA (4 mM final concentration, BioWorld, #40121266-1) and digested chromatin was incubated for 10 min at 65° C shaking. All aliquots from the same experimental conditions were pooled together and centrifuged at 1000x g for 5 min at RT. Fragmented DNA was washed with 500 μl 1X T4 DNA Ligase Reaction Buffer (NEB, #B0202S) diluted in water and 2.5% of the total volume was transferred in a new vial as a control of the MNase digestion level. After the addition of 150 μl of 10 mM Tris-HCl pH 7.4, 25 μl 10% SDS (Invitrogen #15553-035), 25 μl 20 mg/ml Proteinase K (GoldBio, #P-480-1), the control sample was incubated overnight at 65° C shaking.

The remaining sample was centrifuged at 1000x g for 5 min at RT, the pellet was resuspended in 90 μl of freshly prepared Micro-C Master Mix 1 (10 μl T4 DNA Ligase Reaction Buffer (NEB, #B0202S), 68 μl H_2_0, 5 μl 10 U/μl T4 PNK (NEB, #M0201L)) and incubated 15 min at 37° C with 800 rpm shaking. 10 μl of 5 U/μl Klenow Fragment (NEB, #M0206L) were then added to the tube and the suspension was incubated 15 min at 37° C with 800 rpm shaking. T4 PNK enzyme catalyzes the removal of 3’-phosphate and the addition of 5’-phosphate whereas the Klenow enzyme, in the absence of nucleotides, exhibits a 3’-5’ exonuclease activity promoting the removal of the 3’-overhang ends. Subsequently, 50 μl of Micro-C Master Mix 2 (10 μl 1 mm Biotin-dATP (Jenna Bioscience, #NU-835-Bio14-S), 10 μl 1 mm Biotin-dCTP (Jenna Bioscience, #NU-809-BioX-S), 1μl 10 mM mixture of dTTP and dGTP ((10 mM dTTP, NEB, #N0443S), (10 mM dGTP, NEB, #N0442S)), 5 μl 10X T4 DNA Ligase Reaction Buffer (NEB, #B0202S), 0.25 μl 20 mg/ml BSA (NEB, #B9000S), 23.75 μl H_2_0) were added to the fragmented DNA and the reaction was incubated at 25° C for 45 min with shaking. During this step, DNA overhangs are filled with both biotinylated and unbiotinylated nucleotides. The enzymatic reaction was stopped with 9 μl of 0.5 M EDTA (Invitrogen, #15575-038) and incubated at 65° C for 20 min with 800 rpm shaking.

Cells were collected by centrifugation at 1000x g for 5 min RT and the pellet was resuspended in 500 μl of Micro-C Master Mix 4 (422.5 μl H_2_0, 50 μl 10X T4 DNA Ligase Reaction Buffer (NEB, #B0202S), 2.5 μl 20 mg/ml BSA (NEB, #B9000S), 400 U/μl Ta DNA Ligase (NEB, #M0202L)) and incubated at 25° C for 2.5 hours with gentle shaking. After proximity ligation, cells were centrifuged at 1000x g for 5 min RT, resuspended in 200 μl of freshly prepared Micro-C Master Mix 4 (20 μl 10X NEBuffer #1.1 (NEB, # B7001S), 170 μl H_2_0, 10 μl 100 U/L Exonuclease III (NEB, #M0206L)) and incubated at 37° C for 15 min with shaking. To reverse crosslinked the DNA and digest proteins and RNA, 25 μl 10% SDS (Invitrogen #15553-035), 25 μl 20 mg/ml Proteinase K (GoldBio, #P-480-1) were added to the sample, and the cell suspension was incubated overnight at 65° C shaking.

Fragmented DNA was extracted using Ultrapure Phenol Chloroform Isoamyl Alcohol (Invitrogen, #15593-03) and directly incubated with previously prepared Dynabeads MyOne Streptavidin C1 (Invitrogen, #65001).

30 μl of Dynabeads MyOne Streptavidin C1 (Invitrogen, #65001) were washed two times with 300 μl 1x TBW (5 mM Tris-HCl pH 7.4, 0.5 mM EDTA, 1 M NaCl, 0.05% Tween20 (Sigma Aldrich, #P7949)). Beads were then resuspended in 200 μl 2x B&W (10 mM Tris-HCl pH 7.4, 1 mM EDTA, 2 M NaCl). 200 μl of ready streptavidin beads were added to 200 μl of DNA sample and incubated for 20 min at RT with rotation. Beads were then washed twice with 300 μl 1x TBW, once with 100 μl 0.1x TE (1 mM Tris-HCl pH7.4, 0.1 mM EDTA pH 8), and resuspended in 50 μl 0.1x TE. Sequencing libraries were prepared with NEBNext UltraII DNA Library Prep Kit for Illumina (NEB, #E7645L) following the manufacturer’s instructions except for the purification and size selection steps. After the USER enzyme incubation, beads were washed twice with 1x TBW, once with 0.1x TE, and then resuspended in 20 μl of 0.1x TE. To determine the correct number of PCR cycles, 1 μl of streptavidin-biotin DNA underwent 16 amplification cycles. After PCR, DNA concentration was measured, and the DNA fragment size was checked on 1% agarose gel. Once established the proper number of PCR cycles, NEBNext Multiplex Oligos for Illumina (NEB, #E7600S) protocol was followed for PCR amplification and sample indexing. Amplified DNA was then purified with 0.9X SPRIselect size selection beads (Beckman Coulter, #B23319) and eluted in 20 μl 0.1x TE.

#### Target enrichment

Probe panels targeting the locus of interest (chr8 126700640-129900000) were created based on Twist Bioscience requirements. 80-mer probes were designed to cover the entire region without any overlaps. The capture assay was performed following Twist Target Enrichment Standard Hybridization v1 Protocol (Twist Bioscience) and the two biological replicates were handled separately.

For each biological replicate, 4 μg of indexed Micro-C library pool containing all time points were used as starting material for the pulldown. DNA libraries were first concentrated with 1.8X SPRIselect size selection beads (Beckman Coulter, #B23319) and DNA was eluted in 12 μl Twist Universal Blockers (5 μl Blocker Solution and 7 μl Universal Blockers (Twist Bioscience, #100578)). DNA fragments were then incubated together with the biotinylated-probe solution for 16 hours at 70° C. Streptavidin Binding Beads (Twist Bioscience, #104325) were used to pull-down locus-specific hybridized DNA fragments. After several washed, the target library underwent a second round of PCR amplification using Equinox Library Amp Mix (2x) (Twist Bioscience, #104107) and post-amplification library purification with DNA purification Beads (Twist Bioscience, #104325).

Before the sequencing submission, DNA concentration was measured via Qubit 2.0 Fluorometer (Invitrogen), and DNA fragment size distribution was checked via TapeStation (Agilent). Samples were sequenced on an Illumina NextSeq 2000 on 50 base pair-end mode.

#### Data analysis and visualization

RCMC paired-end reads were downloaded as .fastq files and processed with *Distiller* pipeline (https://github.com/mirnylab/distiller-nf). Briefly, read pairs were aligned against the human reference assembly hg38. Aligned reads were parsed, PCR duplicates were removed, and pairs files were generated. Pairs files obtained from the *Distiller* pipeline were used in further analyses.

To retain only those reads that both mates align on the region of interest, a tabular file containing chromosome 8 coordinates was created and pairs files were filtered with *pairtools* select option of *pairtools* package (https://github.com/open2c/pairtools). A 50-bp bin file of the human reference assembly hg38 was used to convert filtered reads into .cool files using cooler *cload* pairs of the cooler package (https://github.com/open2c/cooler). Cool files were then converted into multiresolution cooler files (500bp, 1kb, 2kb, 5kb, 10kb, 50kb, 100kb, 250kb, 500kb, 1Mb) using the cooler *zoomify* option of the cooler package (https://github.com/open2c/cooler).

RCMC data were visualized with *cooltools* package (https://github.com/open2c/cooltools) in *python*, plotted at 10 kb resolution and contact matrixes were exported as .pdf files. Contact matrices were extracted with the *cooltools fetch* function with settings *balance=False and sparse=False* to avoid biases introduced by loci copy number (figure 4 G, L, M). For differential contact matrixes showing the log2 fold change of interactions across the c-MYC TAD, the contacts of each matrix were normalized for read coverage before division.

#### Coverage plots

The CTCF ChIP seq profiles and peaks were downloaded from ENCODE (ENCFF204ZDF and ENCFF502CZS, respectively). The CTCF motif was downloaded from JASPAR (MA0139.1) and converted to *homer* format in R and mapped with homer against the hg38 genome. CTCF motifs within a window of +/-150 bp of a CTCF peak and with a signal strength larger than 75 were used to indicate CTCF directionality (Figure 4 J, K). For MNase-Seq coverage plots, reads collected in the *pairs* file generated by *distiller* were shifted by 73 bp in respect to strand directionality to obtain theoretical nucleosome dyads. To normalize the data across samples, a window of distal of the cleavage site was selected (129.0 Mbp – 129.5 Mbp). The read coverage was computed in 1000 bp windows across the c-Myc TAD and smoothed with a 50 bin role mean (Figure 4 J, K).

#### RNA FISH analysis of c-MYC and PVT1 focal accumulations

RNA FISH was performed for the detection of c-MYC and PVT1 focal accumulations (Fig. 5 and Fig. S5). All reagents were kept RNAse-free and buffers were freshly reconstituted with deionized formamide according to the manufacturer’s recommendations. FISH probes (PVT1 exonic labeled with Quasar 670 dye (Stellaris, BioNordika VSMF-2307-5, MYC exonic labeled with Quasar 570 dye (Stellaris, BioNordika VSMF-2230-5)) were dissolved in TE pH8.0 to a final concentration of 12.5 µM. Cells were grown, fixed and permeabilized using the same protocol as for immunofluorescence stainings. Coverslips were then incubated in buffer A (Stellaris, Nordic Biolabs SMF-WA1-60) for 5 minutes at room temperature. For two coverslips, 1 µl of each FISH probe was mixed with 99 µl hybridization buffer (Stellaris, Nordic Biolabs SMF-HB1-10) and cells were allowed to hybridize overnight in a wetchamber at 37°C. For washing, coverslips were transferred to freshly reconstituted buffer A and incubated at 37°C for 30 minutes, followed by 30 min incubation with buffer A supplemented with 4ʹ,6-diamidino-2-phenylindole dihydrochloride (DAPI, Molecular Probes, D1306, 0.5 μg/ml). After a final wash with buffer B (Stellaris, Nordic Biolabs SMF-WB1-20) for 5 minutes at room temperature, coverslips were mounted using solid mounting medium (Abberior, MM-2013-2X15ML) and allowed to cure overnight. Cells were then imaged using the LSM880 confocal microscope (Zeiss LSM880) and images were analysed using the Imaris Cell Analysis software (version 10.0.0).

### Supplemental figures and legends

**Fig. S1.**
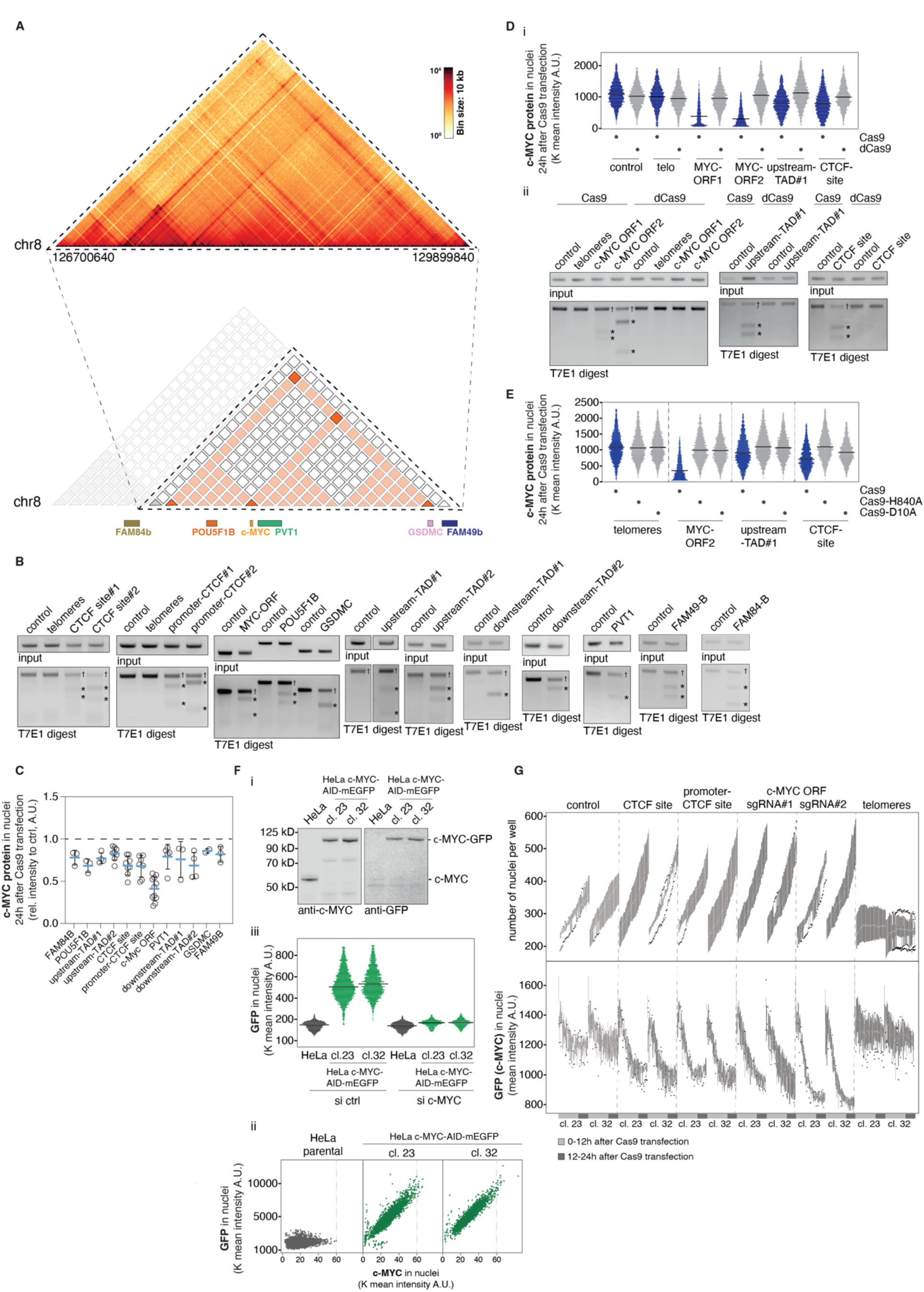
(related to Fig. 1). Cas9 RNP cleavage efficiently targets the entire cell population. (A) Heatmap showing genomic contact frequencies with 10 kb resolution within the c-MYC TAD derived from Region-Capture Micro-C in untreated HeLa cells used throughout this study (top); derived simplified model of the c-MYC TAD (bottom). (B) T7E1 assay to assess the Cas9 cleavage at indicated target sites. EtBr-stained agarose gels showing PCR products over the targeted sites 24h after Cas9 RNP transfections before (upper panels) and after T7E1 nuclease digestion (lower panels). Crosses indicate uncleaved PCR products, asterisks indicate specific T7E1 cleavage fragments. (C) Mean c-MYC protein expression at indicated genomic distances from the c-MYC ORF 24h after Cas9 RNP transfection; each dot represents one independent biological replicate. (D)(i) Mean c-MYC protein expression measured by QIBC in n=3200 cells per sample 24h after transfection with either Cas9 RNPs (blue) or dCas9 RNPs (grey) with sgRNAs targeting indicated sites. (ii) T7E1 assay as in (B) to confirm specific cutting of the indicated target sites by Cas9 and absence of cutting in the same sites in cells transfected with dCas9. Crosses indicate uncleaved PCR products, asterisks indicate specific T7E1 cleavage fragments. (E) Mean c-MYC protein expression measured by QIBC as in (D) in n=3000 cells per sample 24h after transfection with either Cas9 RNPs (blue) or nickase Cas9 RNPs with indicated mutations (grey) and sgRNAs targeting indicated sites. (F) Endogenous tagging of c-MYC with the AID-mEGFP tag in HeLa cells. (i) Analysis of c-MYC-AID-EGFP expression in two selected independent clones compared to the parental cell line obtained by immunoblotting for c-MYC (left) and GFP (right). (ii) QIBC analysis of c-MYC protein levels in parental HeLa cells (grey) and two independent endogenously tagged clones (green) showing high degree of correlation between the c-MYC and the GFP staining. Maximum expression level in the parental cell line is marked by the blue dashed line; n>1500 cells per sample. (iii) QIBC analysis of c-MYC protein levels in parental cells (grey) and the two endogenously tagged clones (green) immunostained for GFP; to confirm specificity, cells were treated with control siRNA (left) or siRNA against c-MYC (right); n>1500 cells per sample. (G) Live imaging of c-MYC protein expression and cell proliferation after Cas9 RNP transfection. Number of cell nuclei (top) and mean expression of c-MYC-AID-EGFP (bottom) in two independent endogenously tagged c-MYC-AID-EGFP clones with n=200-500 cells per condition per timepoint after transfection with indicated Cas9 RNPs.

**Fig. S2.**
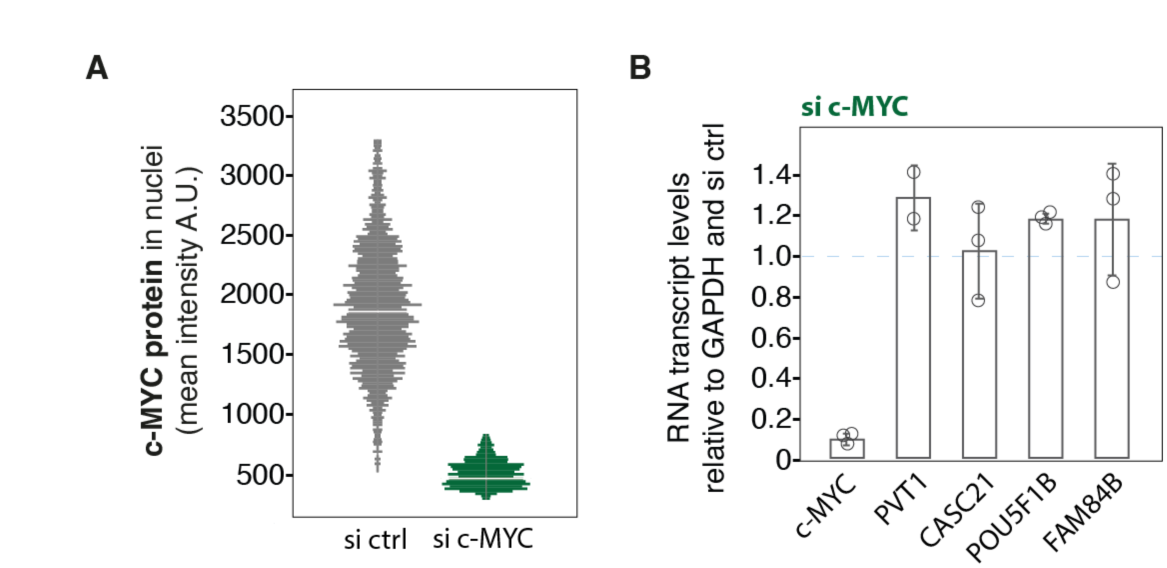
(related to Fig. 2). Local transcript deregulation in the c-MYC TAD subjected to Cas9 cutting is not due to lower c-MYC levels. (A) QIBC control for c-MYC depletion by siRNA, data from a representative experiment from (B); n=1800 cells per condition. (B) HeLa cells treated with control and anti c-MYC siRNA for 24 hours were analysed for transcript levels of c-MYC, PVT1, CASC21, POU5F1B and FAM84B by RT-qPCR as in Fig. 2. Dots show replicates, error bars are standard deviation.

**Fig. S3.**
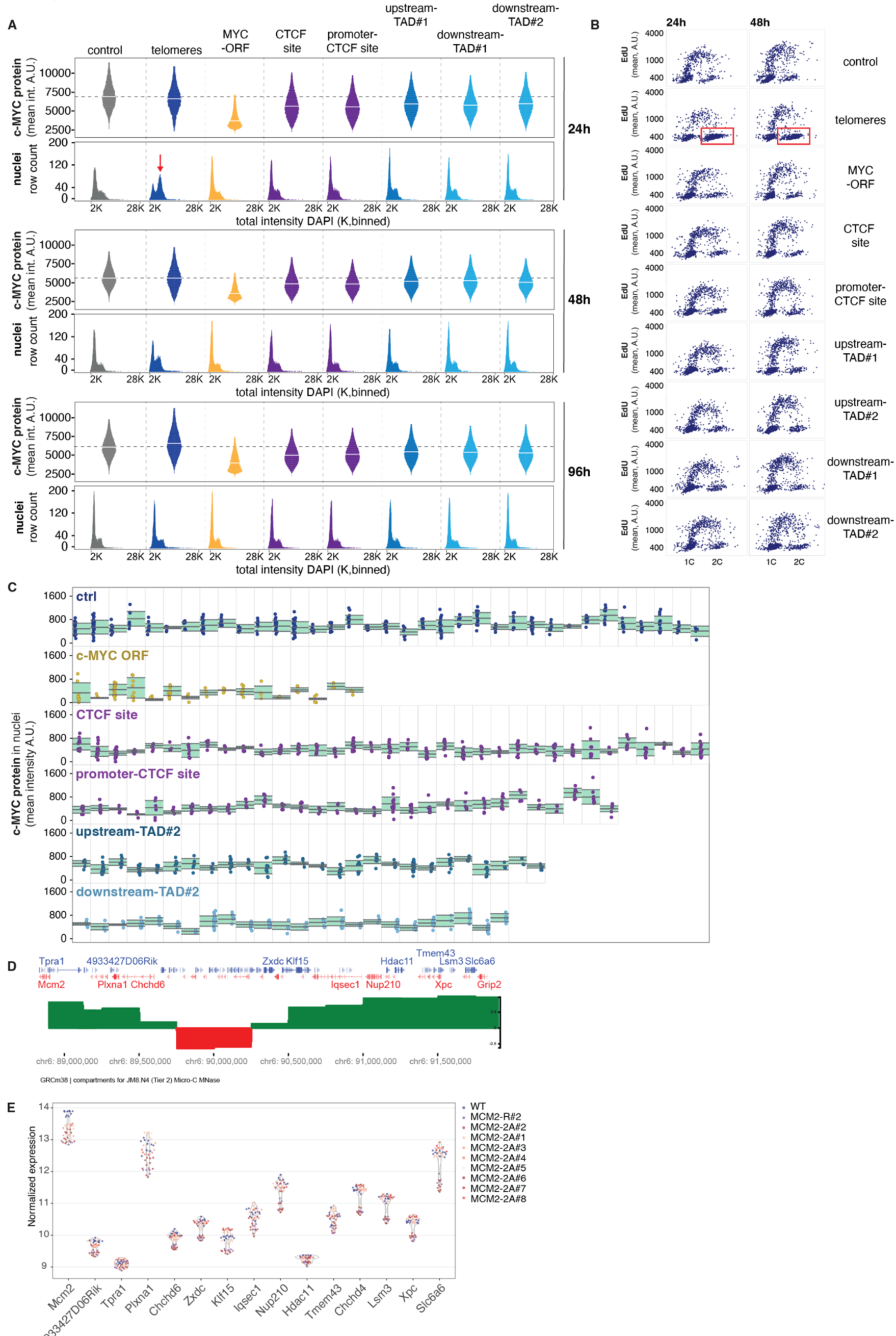
(related to Fig. 3). A single DSB stably alters the transcriptional output across topologically defined genomic loci. (A) Extended data from the experiment shown in Fig. 3B depicting QIBC analysis of mean c-MYC expression at indicated timepoints after Cas9 RNP transfection (upper panels) and the corresponding cell cycle profiles determined by DNA content distribution (lower panels). Red arrow indicates G2 arrest after cutting with a telomeric guide. (B) Cell cycle profiling determined by QIBC analysis of EdU-pulsed cells (n=1000 cells per condition) at indicated timepoints after RNP transfection. Red boxes indicate G2 arrest after cutting with a telomeric guide. (C) Primary QIBC data from the experiment shown in Fig. 3D, E. Dots represent mean c-MYC protein levels in single cells derived from individual clones (separated by vertical lines) after the indicated Cas9 RNP transfections. Green boxes highlight the variance around the median value per clone. (D-E) Genome editing of the *Mcm2* gene causes mild repression within the *Mcm2* locus beyond compartment boundaries. (D) Visualization of compartments signal upstream to *Mcm2* gene calculated using mESC Micro-C data (4DNFILGE5LQU, 4DN portal)(*41*). PC1 analysis of A (green) and B (red) compartments. (E) Dot-plot showing normalized expression of WT, MCM2-2A and MCM2-R cells of selected genes (related to Fig. 3G). Each dot represents the relative expression value of one biological replicate.

**Fig. S4.**
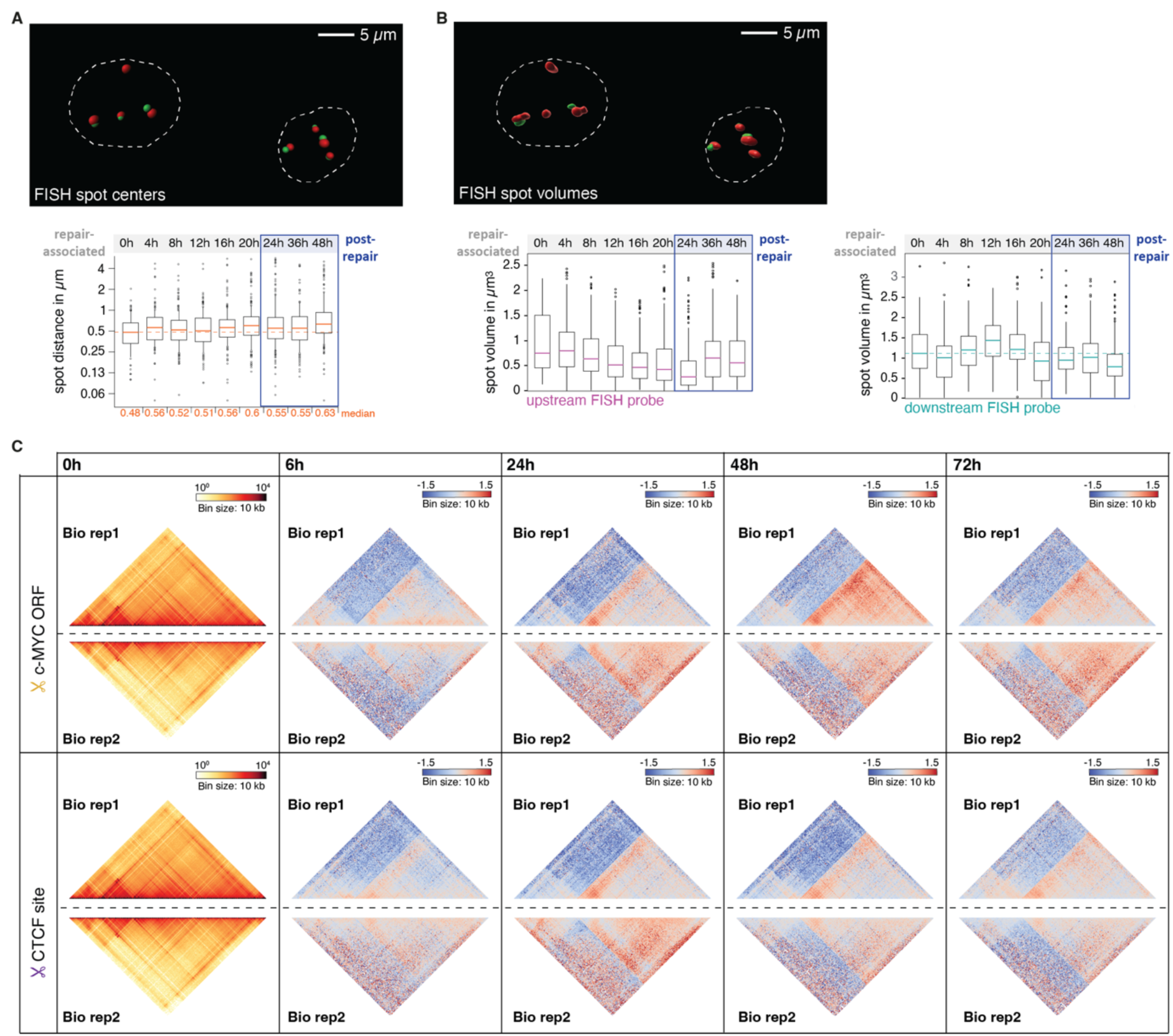
(related to Fig. 4). Extended DNA FISH segmentation and Region-Capture Micro-C analyses to illustrate lasting DSB-induced rearrangements of the c-MYC TAD. (A) Representative image (top) illustrating the segmentation mask applied in Fig. 4C, D and Fig. S4A to calculate the spot centers for each individual FISH probe and subsequently derive the distance between them. Scale bar represents 5 µm. Extended data (bottom) for the experiment shown in Fig. 4C, D; distances between the DNA FISH probes are depicted; horizontal bars mark the median, the dotted line indicates the 0h timepoint, numbers at the bottom are the median values for each timepoint. (B) Representative image (top) illustrating the segmentation mask applied in Fig. 4E, F and Fig. S4B to calculate the spot volumes for each individual FISH spot. Scale bar represents 5 µm. Extended data (bottom) for the experiment shown in Fig. 4E, F; the volumes of the indicated FISH signals in µm_3_ are depicted; horizontal bars mark the median, the dotted line indicates the 0h timepoint. (C) Region Capture Micro-C (RCMC) contact matrixes at 10 kb resolution from both biological replicates (left panel). Fold change contact matrixes of respective time points after Cas9 cleavage at the c-Myc locus over control from both replicates (right panels). All contact maps are binned at 10 kb resolution. Color scale is set to depict the loss of interactions in blue and the gain of interactions in red.

**Fig. S5.**
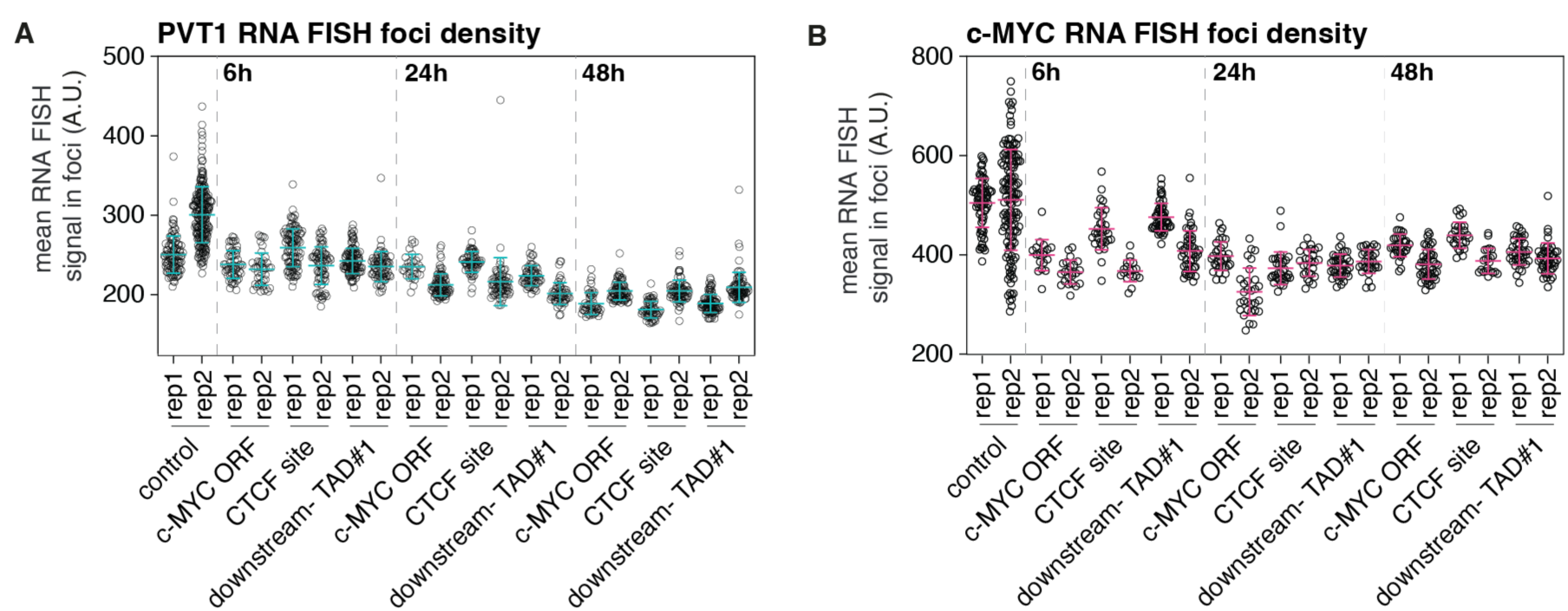
(related to Fig. 5). Extended RNA FISH focus density analysis. Mean intensity of local (A) RNA FISH signal of PVT1 RNA (n=44-227 foci per replicate per condition) and (B) c-MYC RNA (n=20-139 foci per replicate per condition), horizonal bar is the mean of two biological replicates, error bars are standard deviations. Experiments are the same as in Fig. 5 E-F with values for single foci and replicates shown.

**Fig. S6.**
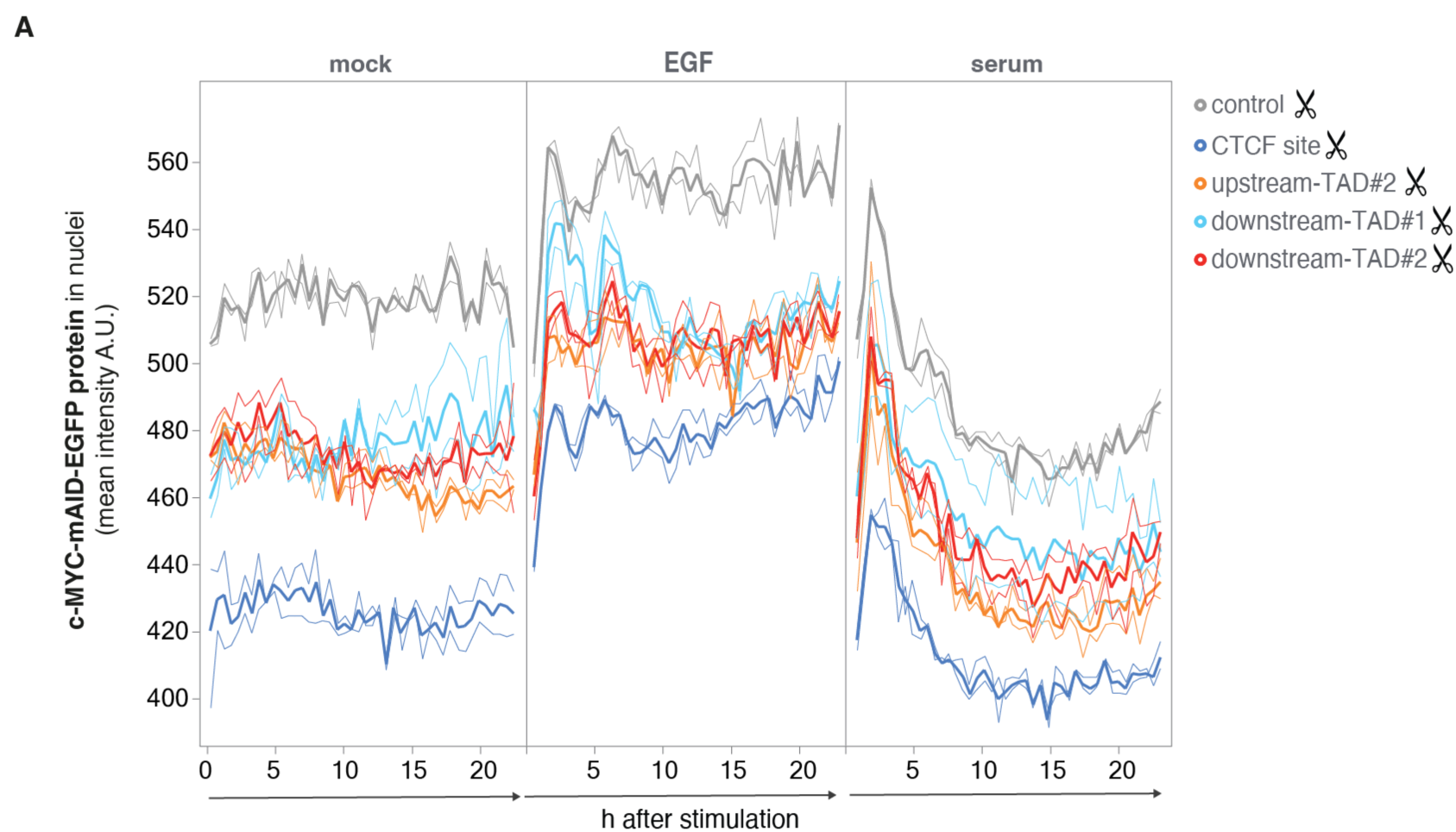
(related to Fig. 6). Attenuated cellular response to EGF stimulation after recovery from a DSB assault in the c-MYC TAD. (A) Primary data for the experiments shown in Fig. 6. The plots depict raw data of mean c-MYC protein levels in two independent endogenously tagged c-MYC-AID-mEGFP HeLa clones after recovery from cutting with the indicated Cas9 RNPs; data are derived from high content live cell imaging without normalization. All cells were serum-starved prior to indicated treatments with 10% FCS, 10 ng/ml EGF and mock, respectively (the complete experimental setup is depicted in Fig. 6B). Bold lines are mean, regular lines are individual clones.

